# Intestinal peroxisomal fatty acid β-oxidation regulates neural serotonin signaling through a feedback mechanism

**DOI:** 10.1101/602649

**Authors:** Aude Bouagnon, Shubhi Srivastava, Oishika Panda, Frank C. Schroeder, Supriya Srinivasan, Kaveh Ashrafi

## Abstract

The ability to coordinate behavioral responses with metabolic status is fundamental to the maintenance of energy homeostasis. In numerous species including *C. elegans* and mammals, neural serotonin signaling regulates a range of food-related behaviors. However, the mechanisms that integrate metabolic information with serotonergic circuits are poorly characterized. Here, we identify metabolic, molecular, and cellular components of a circuit that links peripheral metabolic state to serotonin-regulated behaviors in *C. elegans*. We find that blocking the entry of fatty acyl-CoAs into peroxisomal β-oxidation in the intestine results in blunting of the effects of neural serotonin signaling on feeding and egg-laying behaviors. Comparative genomics and metabolomics revealed that interfering with intestinal peroxisomal β-oxidation results in a modest global transcriptional change but significant changes to the metabolome, including a large number of changes in ascaroside and phospholipid species, some of which affect feeding behavior. We also identify body cavity neurons and an ether-a-go-go related (EAG) potassium channel that functions in these neurons as key cellular components of the circuitry linking peripheral metabolic signals to regulation of neural serotonin signaling. These data raise the possibility that the effects of serotonin on satiety may have their origins in feedback, homeostatic metabolic responses from the periphery.

## Introduction

In both invertebrate and vertebrate species, behaviors such as feeding, movement, reproduction and learning are influenced by nutritional and metabolic signals [1–8]. In mammals, the nervous system actively monitors internal nutritional status by directly sensing specific metabolites like carbohydrates, amino acids and fatty acids, in addition to sensing endocrine signals derived from peripheral tissues [9,10]. These internal nutrient cues are integrated with environmental stimuli and past experiences to orchestrate cohesive and context-appropriate behavioral and physiological responses. Defects in internal metabolic sensing processes contribute to the development of a number of disorders including diabetes, obesity, impaired immune function, neurodegeneration and accelerated aging [8,11–13]. Thus, elucidating the mechanisms by which nutrient status is sensed and communicated between tissues is of critical importance in understanding metabolic homeostasis as well as how metabolism influences myriad physiological and pathophysiological conditions.

Like mammals, *C. elegans* display a range of behavioral and physiological responses to changes in nutrient availability [1,14]. Moreover, as in vertebrate species, the neuromodulator serotonin, 5-hydroxytryptophan, is a key mechanism through which information about food availability is converted to behavioral, physiological, and metabolic responses in *C. elegans* [15–20]. For example, even in the presence of food, worms that lack serotonin display the feeding, egg laying, movement, and metabolic rates that are normally seen when wildtype animals are deprived of food [21]. In contrast, pharmacologic or genetic manipulations that elevate serotonin signaling elicit the range of responses seen when plentiful food supplies are present [19,22,23]. Importantly, serotonin signaling is not simply an *on/off* indicator of food availability but the extent of serotonin signaling allows for animals to fine tune their responses based on their nutritional status and past experiences [3,24,25]. One illustration of this is the effects of varying levels of serotonin signaling on pharyngeal pumping rate, the mechanism by which *C. elegans* ingest nutrients [26,27]. *C. elegans* that have been moved off of their *E. coli* food source reduce their serotonin signaling as well as their pumping rates. Both serotonin signaling and pumping rates are elevated as animals are returned to food [28,29]. However, if animals experience a period of fasting before they are returned to food, they exhibit an even further elevation in feeding rate compared to animals that have only been off of food for brief period of time. The hyper-elevated feeding behavior is accounted for by a correspondingly elevated secretion of serotonin from specific neurons [3]. The hyper-secretion of serotonin and the corresponding hyper-elevated feeding are transient and animals eventually resume the intermediate levels of serotonin signaling and feeding rates seen in well-fed animals [3,29]. Thus, the low, high, and intermediate levels of serotonin signaling correspond to the low, high, and intermediate pharyngeal pumping rates, respectively.

In addition to modulating of food intake behavior, serotonin signaling also affects energy metabolism [30]. *C. elegans* that have been returned to food after a period of fasting transition from a metabolic state that favors energy conservation to an active state of energy utilization. This active metabolic state is driven by elevated serotonin signaling [18,31,32]. If elevated levels of serotonin are maintained by pharmacological or genetic interventions, *C. elegans* exhibit fat loss [33,34]. The effects of serotonin on body fat are not simply a byproduct of its effects on food intake as we and other groups have found that molecular and cellular circuits that link serotonin signaling to peripheral energy metabolism are largely independent from those that regulate feeding [18,22,23,31,32,35]. For example, serotonin secreted from the ADF sensory neurons signals through neurally expressed SER-5 serotonergic receptor to modulate feeding. Yet the SER-5 receptor is not required for the serotonergic regulation of peripheral fat metabolism [18,29]. Instead, serotonin signals through the MOD-1 receptor on URX neurons to promote the release of a neuroendocrine signal that activates triglyceride lipolysis and distinct components of fatty acid oxidation pathways [18,31,32].

In a prior study, we discovered that specific components of lipid oxidation pathways can elicit regulatory effects on feeding behavior [18]. Here, we build upon that finding and describe a feedback mechanism that links peripheral energy metabolism to neuronal serotonin signaling. We find that loss of ACOX-1, a peripheral acyl-CoA oxidase that catalyzes a key step in peroxisomal fat oxidation, affects feeding and egg-laying responses, two serotonin-regulated behaviors. Blunting the utilization of fatty acyl-CoA species, the metabolic substrates of ACOX-1, results in rewiring of peripheral metabolic pathways and ultimately affects the activity of body cavity neurons, which in turn, counteract neural serotonin signaling to influence nutrient-related behaviors.

## RESULTS

### *acox-1* mutants are unresponsive to the feeding stimulatory effects of serotonin

While the serotonergic regulation of fat metabolism is largely distinct from the feeding regulatory pathway, we previously noted two exceptions. RNAi-mediated inactivation of either *acox-1*, encoding a peroxisomal acyl-CoA oxidase, or *cpt-6*, encoding a mitochondrial carnitine palmityoltransferase, not only blocked the fat reducing effects of serotonin but also counteracted the effects of elevated serotonin on feeding [18]. Interestingly, *acox-1* and *cpt-6* function as entry points in the peroxisomal and mitochondrial β-oxidation pathways, respectively [36]. To further investigate how metabolic pathways affect serotonin signaling, we focused on *acox-1* given the availability of a null mutant for this gene at the time that the study was undertaken. Recapitulating our prior RNAi findings, *ad-libitum* fed *acox-1(ok2257)* animals exhibit wildtype feeding rates, yet are unresponsive to the feeding stimulatory effects of exogenous serotonin (Figure 1A, Figure S1A). We previously demonstrated that elevated levels of serotonin signaling exert their effects on fat and feeding pathways by inactivating AMP-activated kinase (AMPK) complexes in distinct neurons [23,29]. As in mammals, the catalytic subunit of AMPK can be encoded by one of two distinct genes, *aak-1* and *aak-2*, in *C. elegans* [37]. Elevated levels of serotonin signaling inactivate the AAK-2 subunit and the hyperactive pumping rate of serotonin-treated wildtype animals is recapitulated by *aak-2* mutants [29]. Loss of *acox-1* suppressed the elevated feeding rates of *aak-2* mutants suggesting that the effects of *acox-1* on feeding are not restricted to exogenously supplied serotonin (Figure 1B).

**Figure 1.**
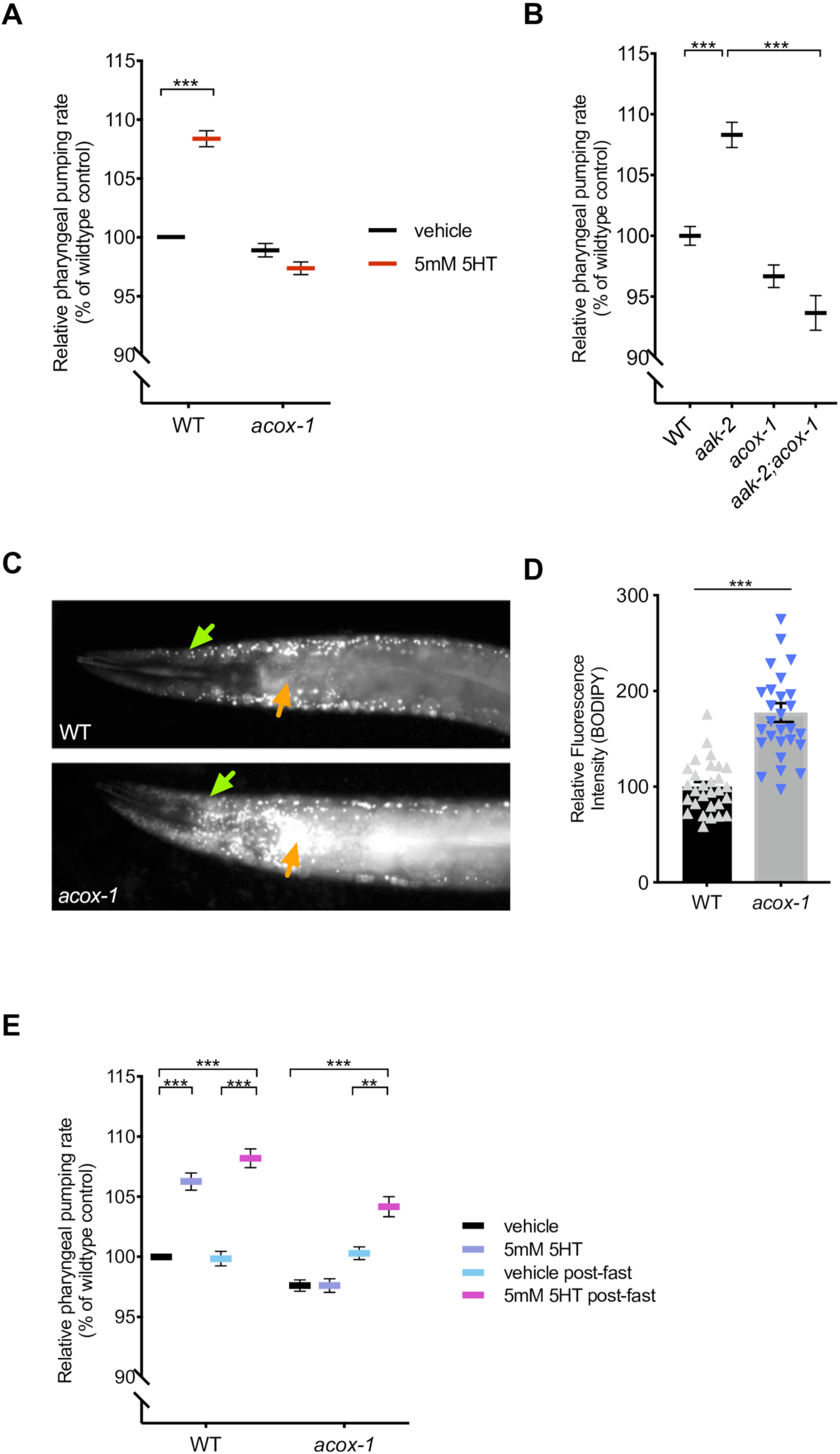
*acox-1* mutants are insensitive to the feeding stimulatory effects of serotonin. **(A)** *acox-1(ok2257)* mutants are resistant to pharyngeal pumping increasing effects of 5mM serotonin. All feeding data is expressed as a percentage of vehicle treated wildtype animals. **(B)** Loss of *acox-1* suppresses the elevated feeding rates of *aak-2* animals. In both **(A)** and **(B)** error bars indicate +/− SEM from mean, n = 50 animals per strain. *** p < 0.0001 ANOVA (Tukey) **(C - D)** *acox-1* mutants accumulate significantly more fat than wildtype animals as assessed by hypodermal (green arrows) and intestinal (orange arrows) BODIPY fluorescence levels. Representative images of BODIPY staining **(C)** and corresponding quantifications of hypodermal BODIPY fluorescence levels **(D)** *** p<0.001, students t-test. **(E)** Fasting *acox-1* mutants for 90 minutes restores their ability to elevate feeding in response to 5mM serotonin. Day 1 adult animals were either fed *ad-libitum* or fasted for 90 minutes than plated on vehicle or 5mM 5HT plates for 60 minutes before assessing pharyngeal pumping rates. Error bars indicate +/− SEM from mean, n = 15 animals per condition. ** p < 0.01, *** p < 0.001 ANOVA (Tukey).

Although the above findings suggested that *acox-1* functions downstream or parallel to serotonin signaling, we sought to rule out the possibility that *acox-1* affects serotonin biosynthesis. Transcription of *tph-1*, the gene that encodes the rate-limiting enzyme of *de novo* serotonin synthesis, is highly dynamic and can be modulated by a range of external and internal cues including food availability, food quality, and stress [24,38–41]. We observed no changes in the transcriptional expression of *tph-1* nor in direct quantifications of serotonin levels in *acox-1* mutants (Figure S1B and S1C). In mammals, defects in peroxisomal fatty acid oxidation pathways are associated with neurodevelopmental disorders due to toxic accumulation of long and very-long chain fatty acids [42]. We had previously shown that elevation of serotonin signaling from the chemosensory, amphid ADF neurons is sufficient to cause elevated pharyngeal pumping [29]. We therefore considered the possibility that the lack of response to serotonin in *acox-1* mutants may be the indirect consequence of a defect in the ADF neurons or other sensory neurons. As one broad examination of neural morphology and development, we used DiI dye staining and found that *acox-1* mutants had properly structured amphid sensory neurons (Figure S1D). Collectively, we found no evidence that deficiencies in serotonin biosynthesis or structural or developmental abnormalities in serotonergic sensory neurons account for the inability of serotonin to elevate feeding rate in *acox-1* mutants.

Based on homology to mammalian acyl-CoA oxidase 1, ACOX-1 is predicted to catalyze the first and rate-limiting step in peroxisomal β-oxidation [43,44]. We used fluorescence intensity of BODIPY labeled fatty acids as a measure of fat accumulation since we and others have previously demonstrated that BODIPY fluorescence corresponds to biochemical and label free methods, such as Coherent anti-Stokes Raman Scattering spectroscopy, measurements of fat content [45,46]. We found that animals lacking ACOX-1 accumulate significantly more fat in their hypodermis and intestines, the two major sites of fat storage, consistent with the notion that *acox-1* mutants have a reduced capacity to break down lipids (Figure 1C and 1D). If the inability of *acox-1* mutants to increase their feeding rate represented a homeostatic response to elevated internal energy stores, we predicted that a period of nutrient depletion should reverse *acox-1* mutants’ feeding behavior. We fasted *acox-1* mutants for 90 minutes, a period of time shown to elicit a coordinated shift in internal metabolic networks towards fat mobilization and energy production, and reintroduced fasted or *ad-libitum* fed animals to either vehicle or serotonin containing plates [47,48]. Fasting restored the ability of *acox-1* mutants to increase their feeding rates in response to exogenous serotonin suggesting that *acox-1* mutants do not simply have a generalized defect in pharyngeal pumping and that a period of starvation can reverse the feeding regulatory effect induced by loss of ACOX-1 activity (Figure 1E). Together, these findings suggest that loss of ACOX-1 activity leads to the generation of a homeostatically regulated signal that can rapidly and reversibly modulate serotoninergic feeding circuits.

### Intestinal ACOX-1 regulates feeding behavior

To elucidate the site of ACOX-1 activity in regulating feeding behavior, we performed tissue-specific rescue experiments. As previously reported, ACOX-1 is expressed in the hypodermis and the intestine, the primary sites of lipid metabolism in *C. elegans* [43] (Figure 2A). Expression of a full length wildtype *acox-1* gDNA sequence in the intestine (via the *vha-6* promoter) but not in the hypodermis (via the *dpy-7* promoter) normalized fat levels in *acox-1* mutants (Figure 2B and 2C). The intestine is the primary metabolic organ in *C. elegans* and carries out numerous metabolic functions including food digestion, nutrient absorption, packaging and secretion [49]. We found that the intestinal expression of wildtype *acox-1* was sufficient to restore the capacity of *acox-1* mutants to elevate feeding rates in response to serotonin treatment (Figure 2D). Thus, intestinal ACOX-1 activity can affect neuronal serotonergic feeding circuits.

**Figure 2.**
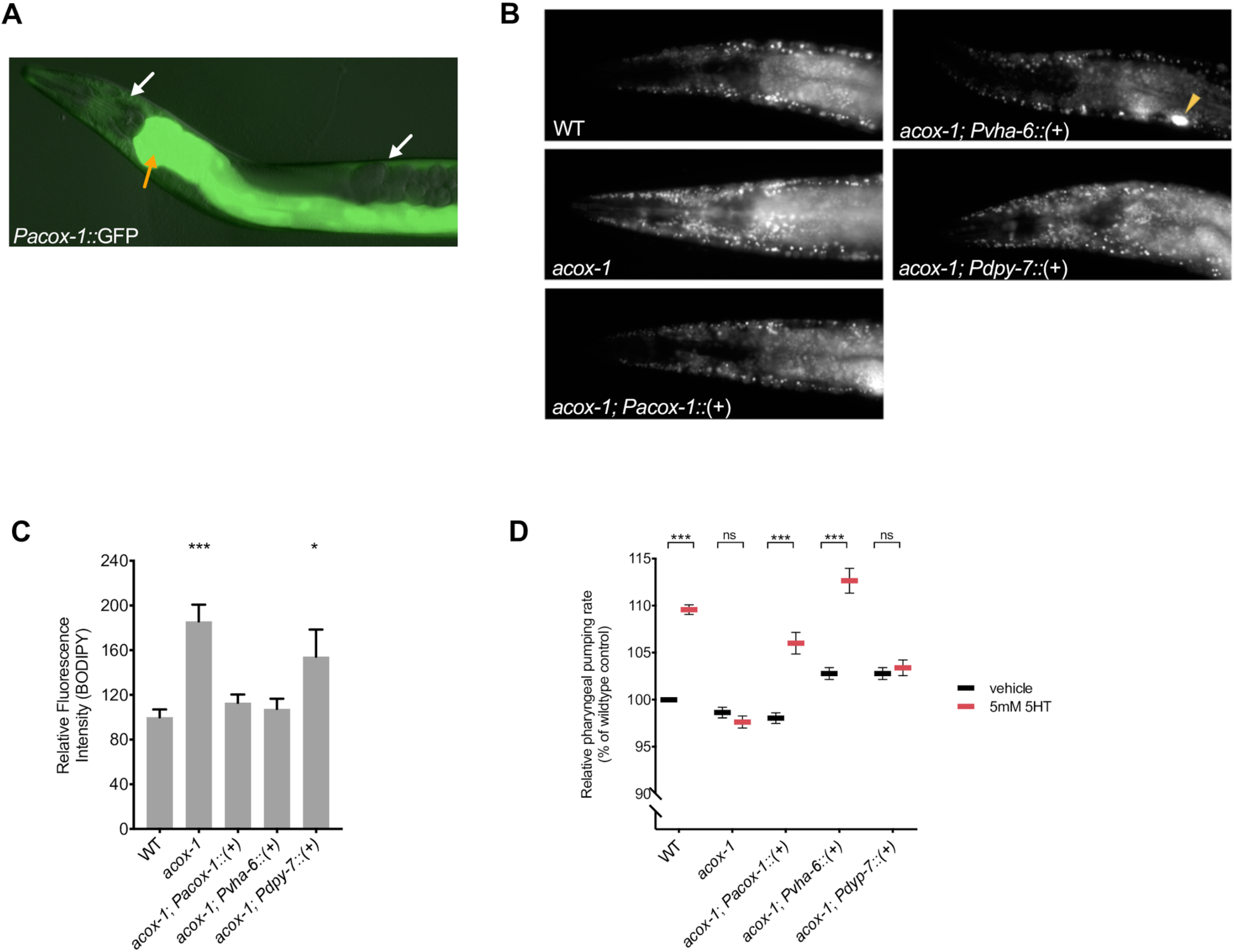
Reconstitution of *acox-1* in the intestine rescues fat and feeding phenotypes. **(A)** *acox-1* is expressed in hypodermal (white arrows) and intestinal (orange arrow) tissues. Merged DIC and green epifluorescent image of a transgenic animal expressing a *acox-1p::gfp* transcriptional reporter. **(B-C)** Reconstitution of *acox-1* gDNA under its own promoter (*Pacox-1*) and under an intestine specific promoter (*Pvha-6*) but not under a hypodermal-specific promoter (*Pdpy-7*) normalizes BODIPY fat levels. Representative images of BODIPY staining **(B)** and corresponding quantifications **(C)** expressed as fluorescence intensity relative to wildtype animals. The yellow arrow in **(B)** indicates a coelomocyte co-injection marker used during transgenic strain generation. Error bars indicate +/− SEM from mean, n>20 per strain. *p<0.05, ***p<0.001, ANOVA (Tukey) **(D)** Reconstitution of *acox-1* gDNA under an intestine specific promoter (*Pvha-6*) restores serotonin responsiveness. Relative pharyngeal pumping rates of indicated strains. Error bars indicate +/− SEM from mean, n > 15 per strain. ***p<0.001, ANOVA (Tukey).

### Modulation of feeding by ACOX-1 requires fatty acyl-CoA synthesis

Acyl-CoA oxidases regulate the rate of metabolic flux through peroxisomes as they govern the first and rate-limiting reaction in peroxisomal metabolic pathways, specifically catalyzing the desaturation of long and very-long chain fatty acyl-CoA esters to 2-trans-enoyl-CoAs [44,50–52]. The *C. elegans* genome encodes seven acyl-CoA oxidases that form homo and heterodimer complexes with distinct substrate specificities [50,53,54]. Structural and biochemical analyses suggest that ACOX-1 homodimers are capable of accommodating a wide-range of fatty acyl-CoA substrates and contribute to peroxisomal β-oxidation of ascaroside lipids [55,56]. Given the enzymatic function of ACOX-1, we hypothesized that its loss leads to an accumulation of fatty acyl-CoA species, which in turn, elicit the anorectic effect. Fatty acyl-CoAs are generated by acyl-CoA synthases (ACS), a family of enzymes that esterify free fatty acids with Co-enzyme A (CoA) [57,58]. To prevent the formation of acyl-CoA thioesters, we acutely exposed animals to Triacsin C, an inhibitor of acyl-CoA synthase activity [59]. This treatment restored the ability of serotonin to cause elevated feeding rate in *acox-1* mutants suggesting that fatty acid esterification is required in order for ACOX-1 to regulate serotonergic feeding cascades (Figure 3A). There at least 20 known or predicted acyl-CoA synthases encoded in the *C. elegans* genome. As a strategy to validate the Triacsin C results and identify the specific synthases involved, we screened through 18 of 20 *acs* genes for which RNAi clones were available. RNAi treatment against multiple *acs* genes, most notably those of *acs-18*, *acs-20* or *acs-22*, suppressed the feeding defects of *acox-1* mutants, suggesting that a degree of redundancy among *C. elegans* acyl-CoA synthases (Figure S3A). As we did not validate the efficacy of each of the 18 RNAis we cannot rule out the possibility that other ACS enzymes also contribute to acyl-CoA pools used by ACOX-1. To further test the notion that fatty acyl-CoAs modulate feeding, we treated animals with oleic acid, a dietary fatty acid, and noted a reduction in pharyngeal pumping rate (Figure 3B). The oleic acid-induced feeding suppression was abrogated in animals that were pretreated with Triacsin C, suggesting that the anorectic effect is induced by oleoyl-CoA or a downstream metabolic derivative.

**Figure 3.**
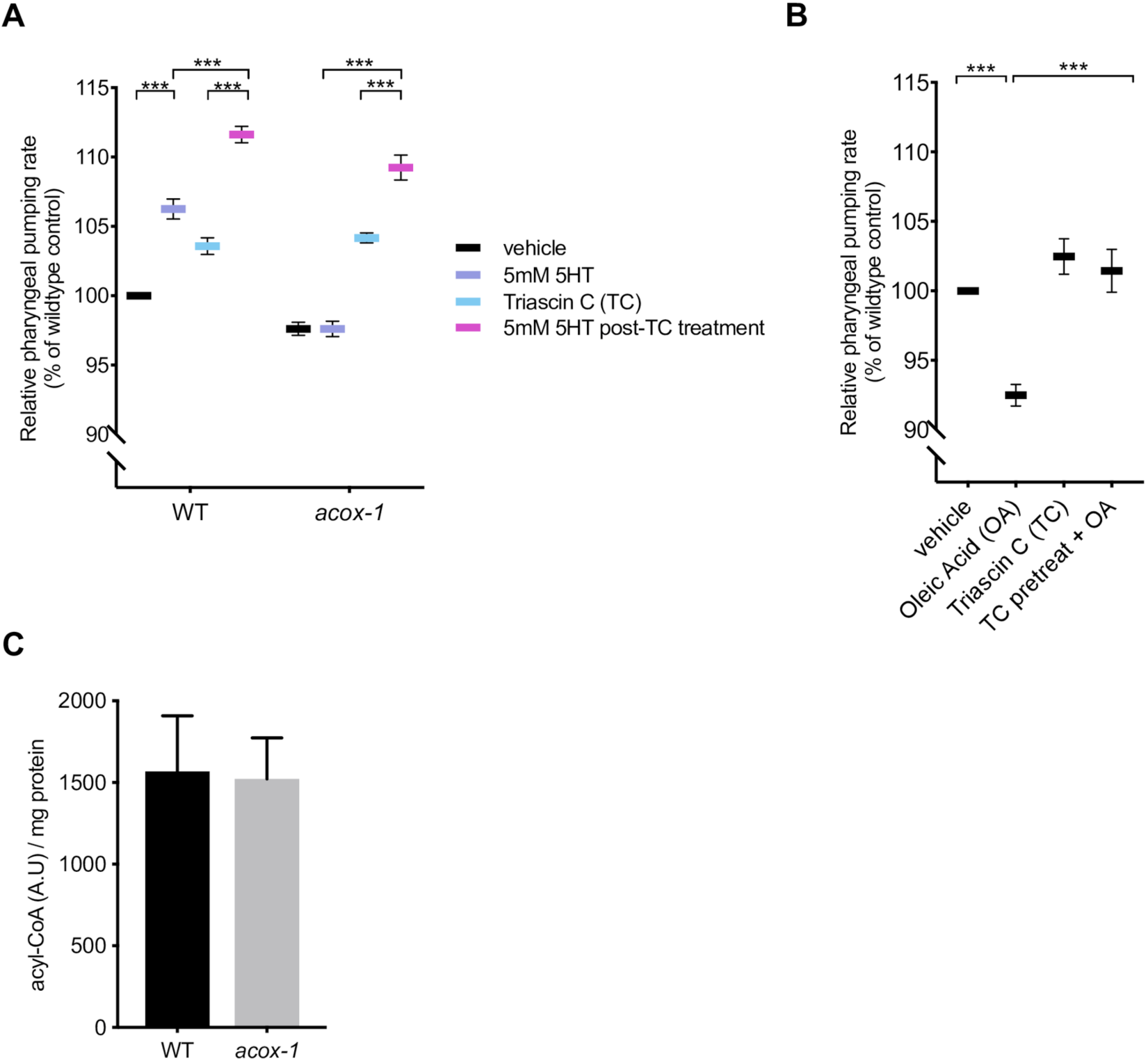
Modulation of feeding by ACOX-1 requires fatty acyl-CoA synthesis. **(A)** Inhibiting acyl-CoA synthesis (ACS) in *acox-1(ok2257)* animals restores the ability to elevate feeding in response to 5mM serotonin. Day 1 adult animals were pre-treated with vehicle or 1µM ACS inhibitor Triacsin C for 90 minutes before being plated on vehicle or 5mM 5HT plates. Feeding was assayed after 60 minutes on assay plates. Error bars indicate +/− SEM from mean, n = 15 animals per condition. ** p < 0.01, *** p < 0.001 ANOVA (Tukey). **(B)** Animals were treated with 1mM oleic acid (OA) or 1µM Triacsin C (TC) for one hour, or pre-treated with 1µM TC or one hour prior to OA treatment before feeding was assayed. Error bars indicate +/− SEM from mean, n > 15 per strain. ***p<0.001, ANOVA (Tukey) **(C)** Acyl-CoA levels in *acox-1* mutants are unchanged relative to WT. Acyl-CoA species were separated and quantified by HPLC and normalized to protein concentrations, n = 3 independent extractions.

To directly evaluate whether loss of *acox-1* causes accumulation of acyl-CoAs, we employed an HPLC-based extraction and detection method from whole animal extracts [60,61]. We found that acyl-CoAs levels were not grossly altered in *acox-1* mutants (Figure 3C). One possible explanation for the absence of elevated acyl-CoA species in *acox-1* mutants is that only a small fraction of total acyl-CoAs in the animals are directed to peroxisomal β-oxidation such that a change in their abundance may not be detectable by our assay. Moreover, acyl-CoA metabolism is known to be highly spatially regulated and acyl-CoA products are selectively synthesized or partitioned in specific tissues [62,63]. Yet another possibility is that acyl-CoAs are substrates for numerous metabolic processes and can be converted into a variety of signaling molecules like ceramides, ascarosides and eicosanoids [54,58,64]. Thus, blocking acyl-CoA utilization by inactivating ACOX-1 may shunt these intermediates into a variety of other metabolic derivatives that ultimately elicit anorectic effect.

### Loss of *acox-1* results in modest transcriptional upregulation of compensatory fat oxidation pathways

To better understand molecular responses to the loss of *acox-1*, we compared the transcriptome *acox-1* mutants to that of wildtype animals using RNA-sequencing. Loss of *acox-1* had a surprisingly limited effect on global gene expression. Our analyses revealed that only 36 out of ∼17,000 genes were differentially expressed in *acox-1* mutants, with only three genes significantly upregulated. Among the differentially expressed genes, the majority were expressed in the intestine, consistent with our finding that this tissue is a major site of action for ACOX-1. Gene Ontology analysis revealed an enrichment for genes involved in “lipid metabolic” and “innate immune” related processes (Table 1). Among the differentially regulated genes, several are predicted to encode for components of peroxisomal β-oxidation including a homolog of acyl-CoA oxidase (ACOX-2), an enoyl-CoA hydratase (ECH*-1.1*) and an ortholog of human bile acid-CoA:amino acid N-acyltransferase (K05B2.4). Two lipases, LIPL-2 and K03H6.2, whose activities are predicted to promote lipid mobilization, were downregulated. Collectively, we interpret these results to mean that the transcriptional responses elicited upon loss of *acox-1* likely compensate for peroxisomal dysfunction by limiting lipid mobilization and by upregulating alternative lipid utilization pathways. Using RNAi, we asked whether inactivation of any of the upregulated genes could counteract the resistance of *acox-1* mutants to serotonin induced feeding elevation but the experiment yielded no such candidates.

**Table 1:**
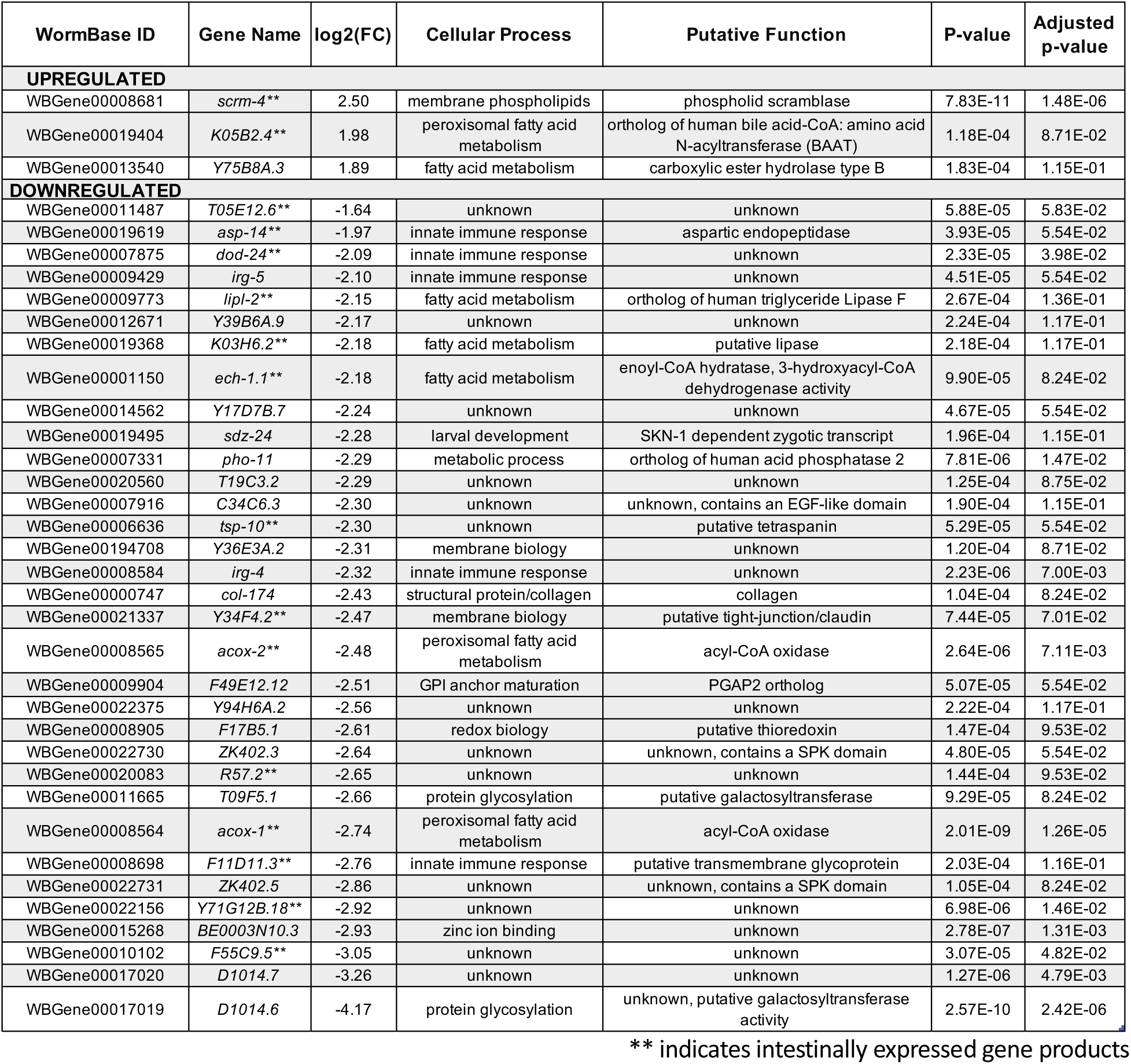
Loss of *acox-1* results in modest transcriptional changes in intestinal and fatty acid metabolic pathways

### Loss of *acox-1* perturbs fatty acid ethanolamide signaling

Next, we sought to obtain a comprehensive overview of the impact of loss of *acox-1* on the *C. elegans* metabolome. For this purpose, we compared the *acox-1* mutant and wildtype metabolomes via untargeted metabolomics using high-pressure liquid-chromatography-high-resolution mass spectrometry (HPLC-HRMS) and the XCMS software platform [65,66]. These analyses revealed that knockout of *acox-1* has a dramatic impact on the *C. elegans* metabolome. Of more than 10,000 significant features detected in the wildtype metabolome, over 500 were at least 3-fold downregulated in *acox-1* mutants. Conversely, we detected more than 500 features that were at least 3-fold upregulated in *acox-1* mutants. To facilitate structural classification of the vast number of detected differential features, we employed molecular networking based on analysis of MS/MS fragmentation patterns [67,68]. These analyses enabled characterization of several metabolite families up- or downregulated in *acox-1* mutants. As expected from previous reports, we found that biosynthesis of most ascaroside pheromones with fatty acid side chains shorter than 9 carbons is abolished or strongly reduced in *acox-1* mutants, whereas abundances of ascarosides with saturated side chains of 9 to 15 carbons are 10- to 50-fold increased [55,56]. In addition, a large number of diacylglycerophosphoethanolamines (DAGPEs), primarily derived from saturated and mono-unsaturated fatty acids with 14 to 18 carbons, were reduced 5- to 20-fold in *acox-1* mutants. Among the metabolites most strongly upregulated in *acox-1* mutants, thiamine and several thiamine derivatives were most prominent, alongside ascr#18 and ascr#22, ascarosides with 11 and 13 carbon sidechains, respectively (Figure 4, Supplemental Table 1).

**Figure 4.**
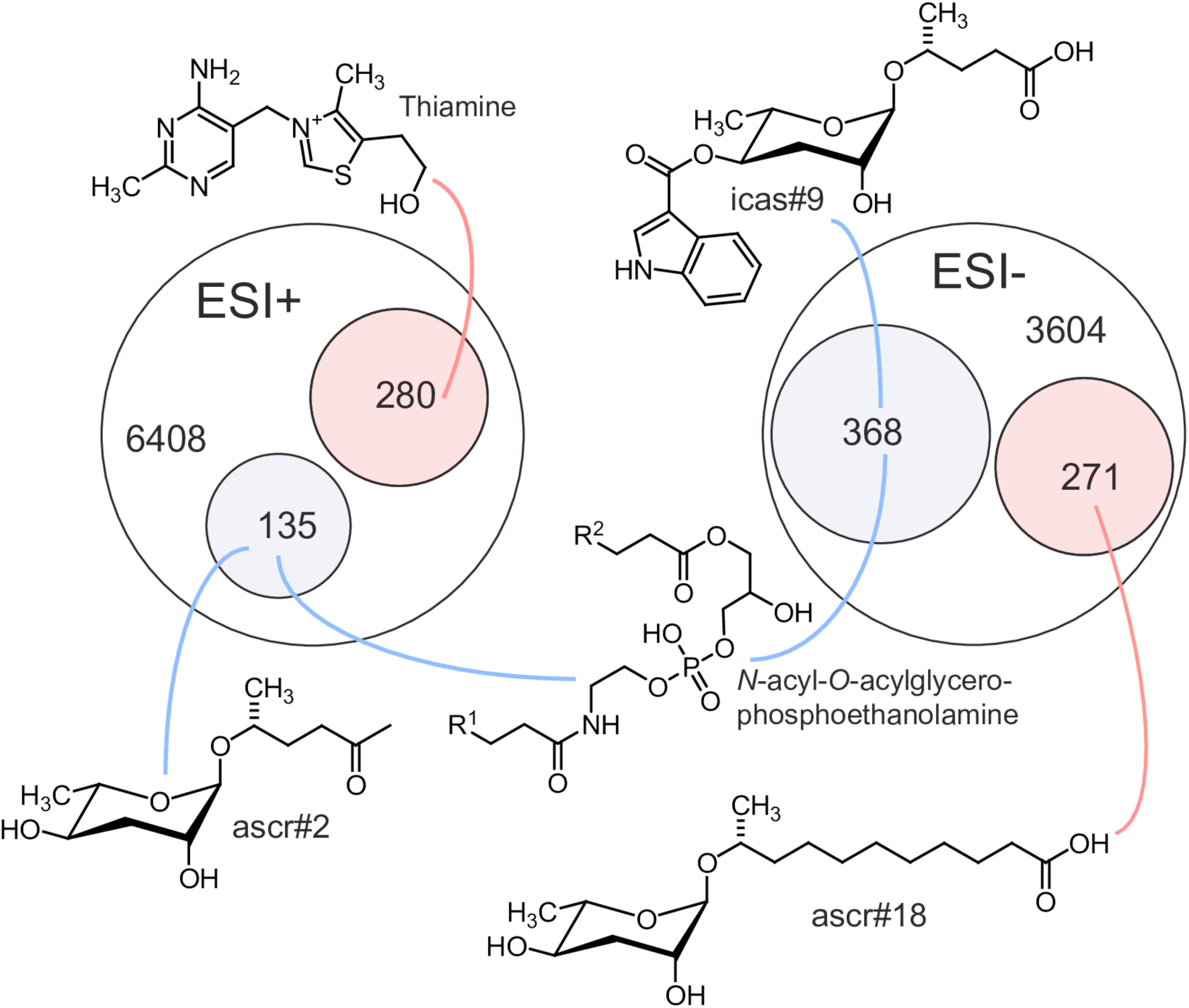
Loss of *acox-1* results large-scale changes to the global metabolome. Venn diagrams showing total numbers of detected features (6408 and 3604) and numbers of metabolites more than three-fold upregulated (pink) or downregulated (blue) in LC-HRMS using positive-ion (ESI+) and negative-ion (ESI−) electrospray ionization in *acox-1* mutants. Though most differentially detected features were unidentifiable, the chemical structures shown here represent examples of the most significantly up or down regulated compounds in identifiable metabolic classes. See Supplemental Table 1.

The very large number of metabolites affected by loss of *acox-1* prevented a comprehensive examination of each of the metabolites. Nevertheless, we used a combination of chemical, genetic, and metabolite add-back experiments to broadly investigate the noted metabolomics changes. We began by considering the possibility that perturbations in ascaroside biosynthesis underlie the feeding defect observed in *acox-1* mutants. Originally identified as constituents of the dauer pheromone, ascarosides are large class of excreted small molecules that regulate development and behavior and whose synthesis can be influenced by metabolic status [69–71]. The acyl-CoA thiolase, DAF-22, catalyzes the terminal step in peroxisomal β-oxidation and plays an essential role in shortening the fatty acid-like-side chains of ascarosides [72]. Though *daf-22* mutants lack ascarosides they still exhibit wildtype feeding rates and are still responsive the feeding increasing effects of serotonin, suggesting that ascaroside biosynthesis and serotonergic feeding regulation are independent of one another (Figure S4A). Though a number of ascaroside species were strongly reduced in *acox-1* mutants, we detected a nearly 30-fold accumulation of ascaroside #18 (ascr#18). To determine if an accumulation of ascr #18 underlies the suppressed feeding response of *acox-1* mutants, we treated wildtype animals with ascr#18 and serotonin. Surprisingly, rather than suppressing feeding, ascr#18 dramatically increased pharyngeal pumping rates of wildtype animals in a manner that was additive with serotonin (Figure S4B). Together, these results suggest that while certain ascarosides species can have feeding regulatory effects, they are unlikely to underlie the specific feeding responses elicited in *acox-1* mutants.

Thiamine and thiamine derivatives were also significantly accumulated in *acox-1* mutants. We next asked if thiamine supplementation was sufficient to block the feeding enhancing effects of serotonin, though did not find this to be the case (Figure S4C). As in mammals, *C. elegans* does not produce thiamine and obtains this essential cofactor from its diet [73]. Given the pervasive changes to *C. elegans* metabolism upon loss of *acox-1*, we speculate that thiamine accumulation in these mutants reflects a general downregulation of enzyme activities that use thiamine derivatives as cofactors.

We next turned our attention to the finding that loss of *acox-1* perturbs the biosynthesis of ethanolamine containing lipid species. Most prominently, we find that diacylglycerophosphoethanolamine (DAGPEs) synthesis is strongly reduced in *acox-1* mutants. DAGPEs likely represent intermediates in the biosynthesis of *N*-acylethanolamines (NAEs), a diverse family of signaling molecules with range of biological roles including in nutrient sensing and mammalian appetite regulation [74–78]. This family of lipids includes the mammalian orexigenic endocannabinoid arachidonoyl ethanolamide (AEA) and anorectic factor oleoyl ethanolamide (OEA) [79]. To test if a reduction in NAEs contributes to ACOX-1 mediated feeding regulation, we inactivated fatty acid amide hydrolase (*faah-1*), an enzyme involved in the hydrolytic degradation of NAEs. We found that inhibition of FAAH-1, which was shown to increase endogenous NAE abundance [80], stimulates pharyngeal pumping rates of both wildtype and *acox-1* mutants (Figure S4D). Thus, in *C. elegans* as in mammals, NAE signaling regulates feeding behavior. As the feeding increasing effects of increasing NAEs were not restricted to *acox-1* mutants, we cannot definitively know that reduced NAE synthesis accounts for the inability of *acox-1* mutants to respond to serotonin. Though this data is intriguing, it is also possible that an as of yet unidentified metabolite or a complex combination of metabolites blunt serotonergic modulation of feeding in *acox-1* mutants.

### ACOX-1 mediated regulation of serotonergic feeding circuits requires EGL-2 activity

We next sought to identify the neural mechanisms that link changes elicited by loss of *acox-1* to serotonergic mechanisms of feeding. The clue that guided us towards an answer emerged unexpectedly by following another phenotype that we had noted in *acox-1* mutants. Relative to wildtype animals, *acox-*1 mutants hold more eggs *in utero* and lay embryos at a later developmental stage (Figure 5A, Figure S4A). This was intriguing since serotonin is also a key modulator of egg-laying behavior [81]. Serotonin controls the excitability of the egg-laying neuromuscular circuit and governs the activity and timing of egg-laying in response to various sensory cues [15–17,82–87]. We found that *acox-1* mutants were markedly less responsive to exogenous serotonin and resistant to the effects of fluoxetine, a serotonin reuptake inhibitor consistent with a reduced response to the excitatory effects of serotonin at the level of the neuromuscular junction (Figure 5C, Figure S4B). As in the context of feeding, reconstitution of *acox-1* in the intestine but not the hypodermis also rescued egg-laying defects of these mutants (Figure 5D and 5E).

**Figure 5.**
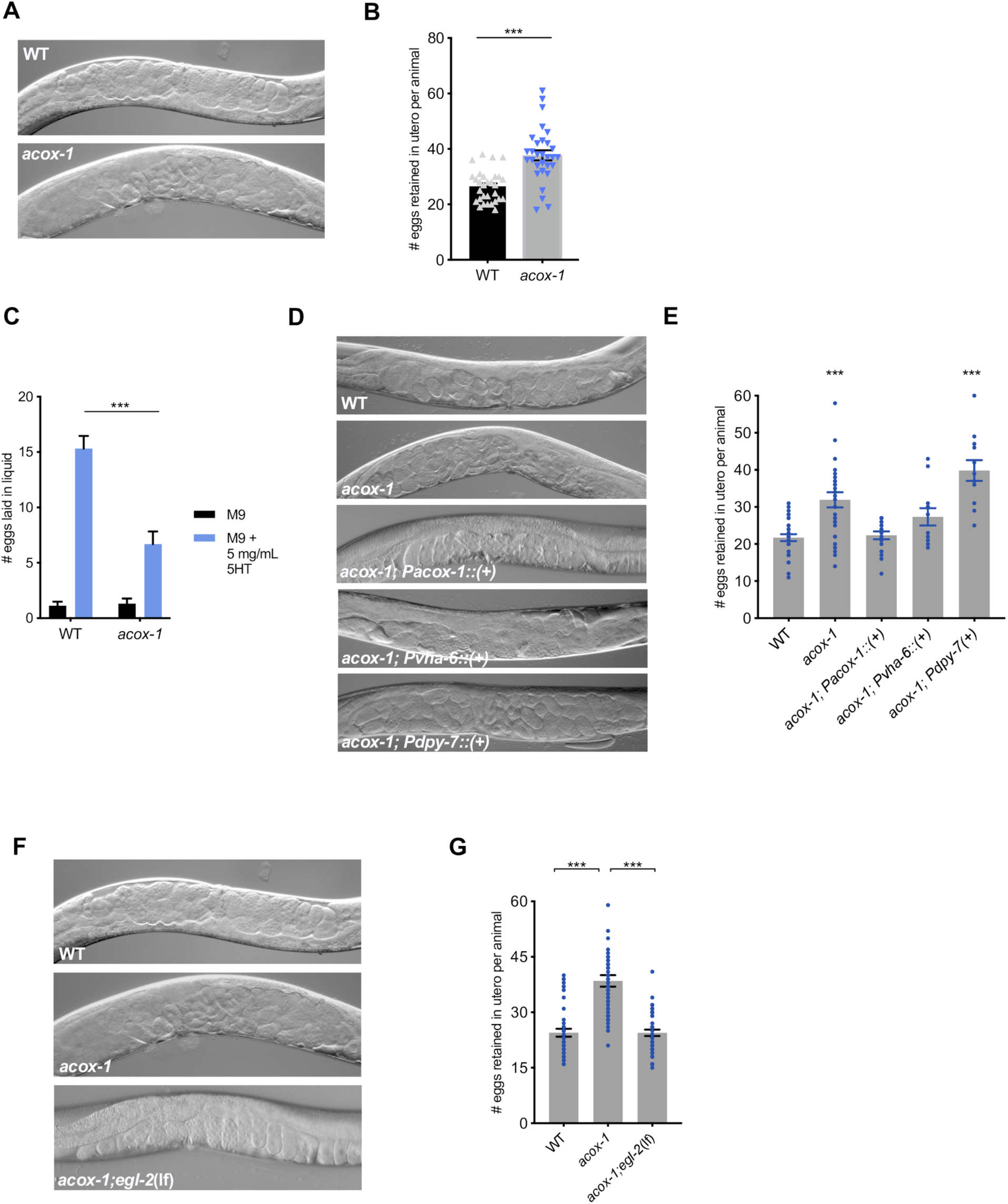
*acox-1* mutants exhibit egg-laying defects. **(A-B)** *acox-1*(ok2257) mutants accumulate significantly more eggs *in utero* than wildtype animals. Representative DIC images **(A)** and quantification **(B)** of eggs retained *in utero*. Error bars indicate +/− SEM from mean, n = 30 animals per genotype. *** p < 0.001 unpaired student’s t-test. **(C)** *acox-1* animals are less responsive to the egg-laying inducing effects of serotonin. Egg-laying response of wildtype and *acox-1* in control buffer (M9) or 5mg/mL serotonin in M9 buffer. Data represents the number of eggs released per animal after a 20-minute exposure to vehicle or drug. Error bars represent +/− SEM from mean. n = 20 animals per condition, *** p < 0.001 ANOVA (Sidak) **(D-E)** Reconstitution of *acox-1* gDNA under an intestine specific promoter (*Pvha-6*) but not a hypodermal specific promoter (*Pdpy-7*) restores egg-laying capacity to *acox-1* mutants. Representative DIC images **(D)** and quantification **(E)** of eggs retained *in utero*. Error bars indicate +/− SEM from mean, n = 30 animals per genotype, *** p < 0.001 unpaired student’s t-test, against a wildtype control. **(F-G)** Loss of *egl-2* rescues *acox-1* egg-laying defects. Representative DIC images of each genotype **(F)** and quantification **(G)** of eggs retained *in utero*. Error bars indicate +/− SEM from mean, n = 30 animals per genotype. *** p < 0.001 unpaired student’s t-test.

The egg-laying neuromuscular circuit has been extensively studied in *C. elegans* and it is well established that egg-laying behavior is strongly influenced by potassium channel activity [88]. Potassium channels are a highly diverse and evolutionarily conserved family of proteins that modulate cellular excitability by regulating the flow of potassium ions (K+) across cellular membranes [89,90]. Gain-of-function mutations in distinct potassium channels have been shown to reduce the excitability of neurons or vulval muscles, and in turn causing egg-laying defects [91–94]. We therefore considered the possibility that aberrant potassium channel activity contributes to the egg-laying dysfunction in *acox-1* mutants. We conducted an RNAi screen of potassium channels with documented roles in egg-laying and assessed their capacity to rescue the egg-laying defect in *acox-1* mutants. RNAi knockdown of the EGL-2 potassium channel normalized egg-laying responses in *acox-1* mutants without influencing baseline egg-laying rates (Figure S4C and S4D). To validate our RNAi results, we crossed *acox-1* mutants with *egl-2*(lf) mutants and found that double mutants resembled wildtype animals in their egg-laying responses (FIGURE 5F and 5G). *egl-2* encodes an ether-a-go-go (EAG) voltage-gated potassium channel that has been shown to modulate the excitability of neuromuscular circuits in response to starvation states, suggesting they may more generally serve as a link between internal nutrient status and neuronal activity [95–97].

To determine if EGL-2 plays a role in ACOX-1 mediated feeding regulation, we took advantage of gain-of-function mutations in *egl-2.* These mutations cause a negative shift the voltage-dependence of these channels and cause pleotropic sensory and behavioral defects by reducing the excitatory capacity of cells in which they are expressed [98,99]. It has been previously reported that *egl-2*(gf) mutants are resistant to the egg-laying inducing effects of serotonin [94,100]. We find that *egl-2*(gf) mutants are also resistant to the feeding enhancing effects of serotonin and thus mimic *acox-1* mutants in their blunted responses to the excitatory effects of serotonin signaling in the context of both egg-laying and feeding (Figure 6A). Both the egg-laying and pharyngeal pumping phenotypes of *egl-*2(gf) mutants can be fully suppressed by *egl-2* RNAi knockdown suggesting that the behavioral effects of aberrant EGL-2 activity can be normalized by inactivating the channel (Figure 6A, Figure S5A). Similarly, RNAi inactivation of *egl-2* restores the ability of *acox-1* mutants to elevate their pumping rates upon exposure to serotonin signaling but without affecting the basal pumping rate in *acox-1* mutants (Figure 6B). Validating the RNAi results, we find that double *acox-1;egl-2(*lf) mutants exhibit wildtype feeding responses to exogenous serotonin (Figure 6C). Finally, we find that the feeding reducing effects of oleic acid also require EGL-2 (Figure 6D). Together, these results suggest that an EGL-2 containing circuit can serve as a regulatory link between metabolic status and feeding behavior.

**Figure 6.**
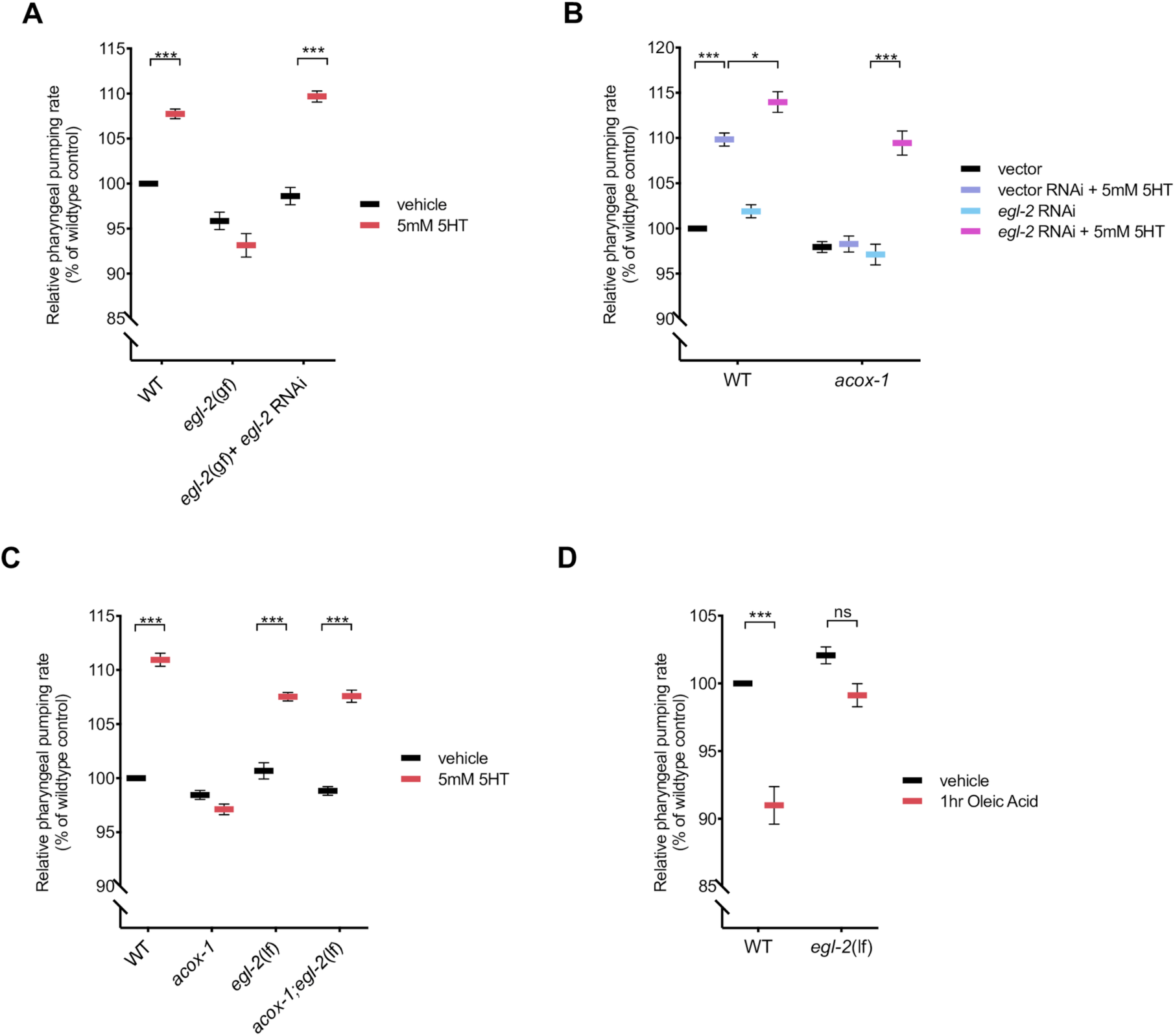
ACOX-1-mediated regulation of serotonergic feeding responses requires the EGL-2 K+ channels. **(A)** Pharyngeal pumping rates of wildtype, *egl-2*(n698), and *egl-2* RNAi treated *egl-2*(n698) mutants on vehicle or 5mM 5HT containing plates. **(B)** Pharyngeal pumping rates of wildtype and *acox-1(*ok2257) animals treated with vector RNAi or *egl-2* RNAi and vehicle or 5mM 5HT **(C)** Pharyngeal pumping rates of indicated strains treated with vehicle or 5mM 5HT. Error bars indicate +/− SEM from mean, n=20 animals per condition. *p<0.05, *** p < 0.001, ANOVA (Tukey) **(D)** EGL-2 is required for animals to reduce feeding in response to oleic acid. Animals were exposed to 1mM Oleic Acid for one hour before feeding was assayed. n> 15 per strain, Error bars indicate +/− SEM from mean, ***p<0.001, ANOVA (Tukey). All feeding data are expressed as a percentage of vehicle treated wildtype animals.

### *acox-1* regulates URX body cavity neuron activity to limit feeding responses

Our results with *egl-2* provided us the opportunity to pinpoint neurons that may serve as a link between peripheral metabolic signals and processes regulated by serotonin signaling. In hermaphrodites, *egl-2* is expressed in a limited subset of sensory neurons including AFD, ALN, AQR, ASE, AWC, BAG, IL2, PLN, PQR and URX [98]. We were intrigued by the expression of *egl-2* channels in body cavity neurons (AQR, PQR and URX), as these are the only neurons with dendritic projections within the pseudocoelom, the rudimentary circulatory fluid utilized by *C. elegans* (Figure 7A) [101]. Given their unique anatomic position, these neurons are hypothesized to mediate bidirectional communication between nervous system and peripheral tissues as they have the capacity to both release and detect circulating signals. Interestingly, neural serotonin signaling is thought to promote intestinal fat metabolism through the release of an neuroendocrine signal from URX neurons [31,32] and fluctuations in peripheral fat metabolism modulate the tonic activity of URX [102]. We find that targeted expression of *egl-2(*gf) in only the body cavity neurons is sufficient to block the feeding increasing effects of serotonin and mimic the effects of loss of *acox-1* (Figure 7B).

**Figure 7.**
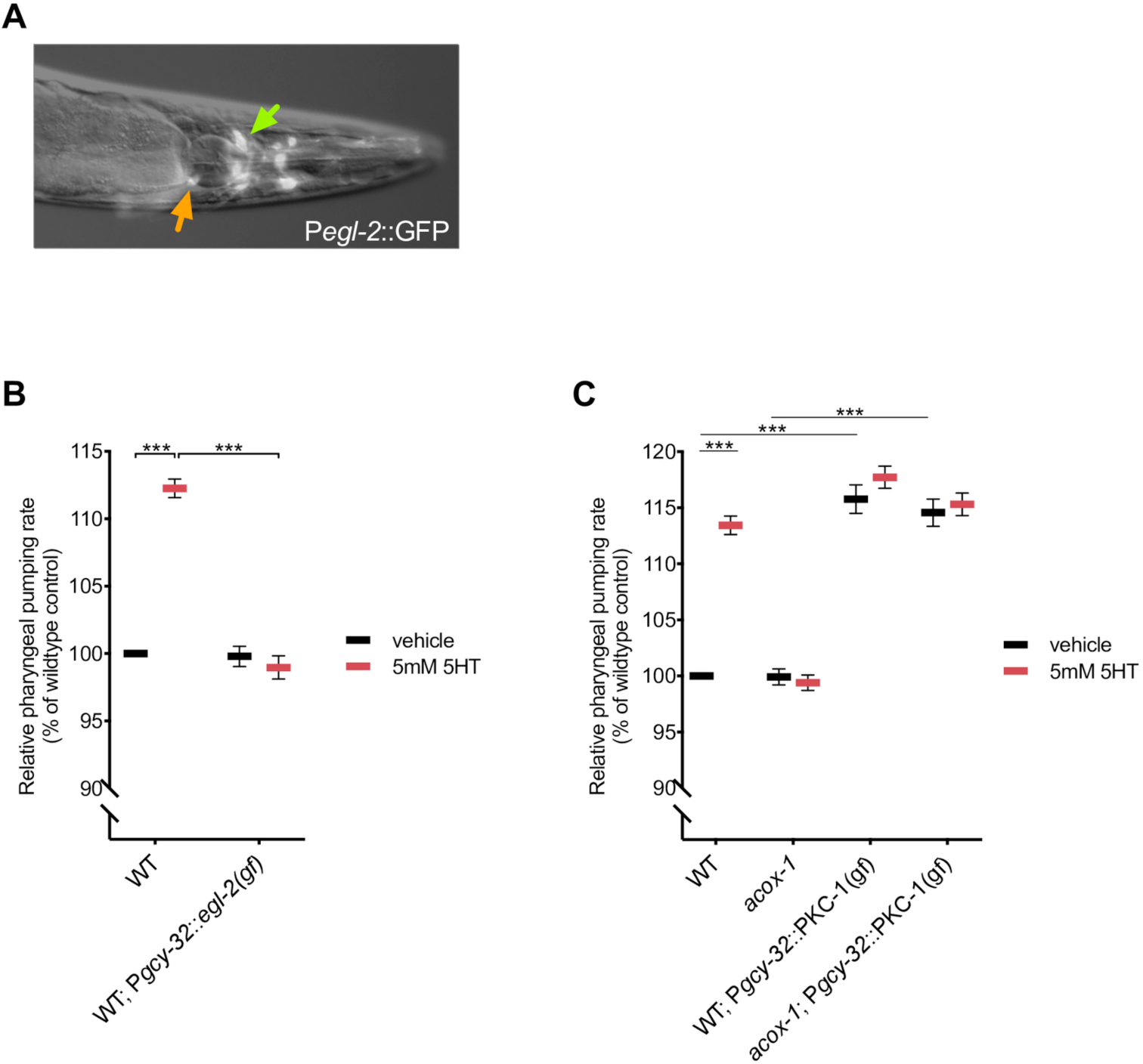
Body cavity neurons modulate feeding behavior. **(A)** EGL-2 is expressed in a limited subset of sensory neurons. Epifluorescent image of a transgenic animal expressing *egl-2p::gfp* transcriptional reporter. Yellow and green arrows indicate the AQR and URX body cavity neurons, respectively. **(B)** Animals expressing *egl-2*(gf) in body cavity neurons do not elevate feeding in response to 5mM 5HT. Pharyngeal pumping rates of wildtype and *pgcy-32::egl-2*(gf) expressing animals treated vehicle or 5mM 5HT. **(C)** Constitutive activation of synaptic release from body cavity neurons stimulates feeding. Pharyngeal pumping rates of wildtype, *acox-1*(ok2257), CX10386 *pgcy-32::pkc-1*(gf) and *acox-1;* CX10386 *pgcy-32::pkc-1*(gf) animals on vehicle or 5mM 5-HT. In **(B)** and **(C)** data are expressed as a percentage of vehicle treated wildtype animals. Error bars indicate +/− SEM from mean, n=15 animals per condition. *** p < 0.001 two way ANOVA (Tukey)

To our knowledge, body cavity neurons have not previously been implicated in the regulation of feeding behavior. To further study their role, we examined the effect of prolonged activation of body cavity neurons on pharyngeal pumping. Activation of PKC-1 has been shown to promote synaptic transmission and neuropeptide release from expressing neurons [103]. We examined the feeding responses of wildtype and *acox-1* mutants expressing constitutively active protein kinase C [PKC-1(gf)] in body cavity neurons and found that this manipulation strongly stimulates pharyngeal pumping in both wildtype and *acox-1* mutants. Importantly, this feeding enhancement could not be further elevated by addition of serotonin, suggesting a common feeding regulatory circuit. (Figure 7C). As genetic inhibition of body cavity neurons by *egl-2(*gf) did not modulate baseline feeding rates, our data suggests that these neurons likely fine-tune feeding responses in distinct conditions rather than governing basal feeding behavior.

To directly assess the effects of ACOX-1 activity on body cavity neuron function, we measured intracellular Ca^2+^ transients from the URX body cavity neurons using the genetically-encoded calcium reporter GCaMP5K [102,104]. We selected URX neurons as calcium dynamics from these cells have been well documented and intriguingly their activity has recently been shown to be modulated by internal metabolic cues and starvation [102,105–107]. URX neurons play a well-documented role in oxygen sensing and are robustly and rapidly activated under atmospheric conditions (21% oxygen) (Figure 8A and 8C) [105,108–110]. To determine if loss of ACOX-1 modulates URX activity, we imaged O_2_-evoked calcium transients in URX neurons in wildtype and *acox-1* mutants. We observed no significant difference in URX responses in wildtype and *acox-1* mutants at 10% oxygen, a concentration at which the tonic URX neurons are held in the “off” state [105]. This result suggests that ACOX-1 does not modulate basal activity of URX (Figure 8B and 8F). However, whereas URX neurons in wildtype animals robustly activate at 21% oxygen as previously documented, O_2_-evoked calcium transients were dramatically inhibited in *acox-1* mutants (Figure 8D and 8D). This reduction in maximal activation (FΔ/F_o_) could not be attributed to altered promoter activity or drift in expression across the tested lines as the level of a co-expressed *flp-8*::mCherry reporter was not measurably different *acox-1* mutants (Figure 8E and 8G). Together, this data suggest that loss of ACOX-1 decreases the sensitivity of URX neurons to excitatory stimuli, supporting the model that body cavity neuron activity is suppressed in *acox-1* mutants.

**Figure 8.**
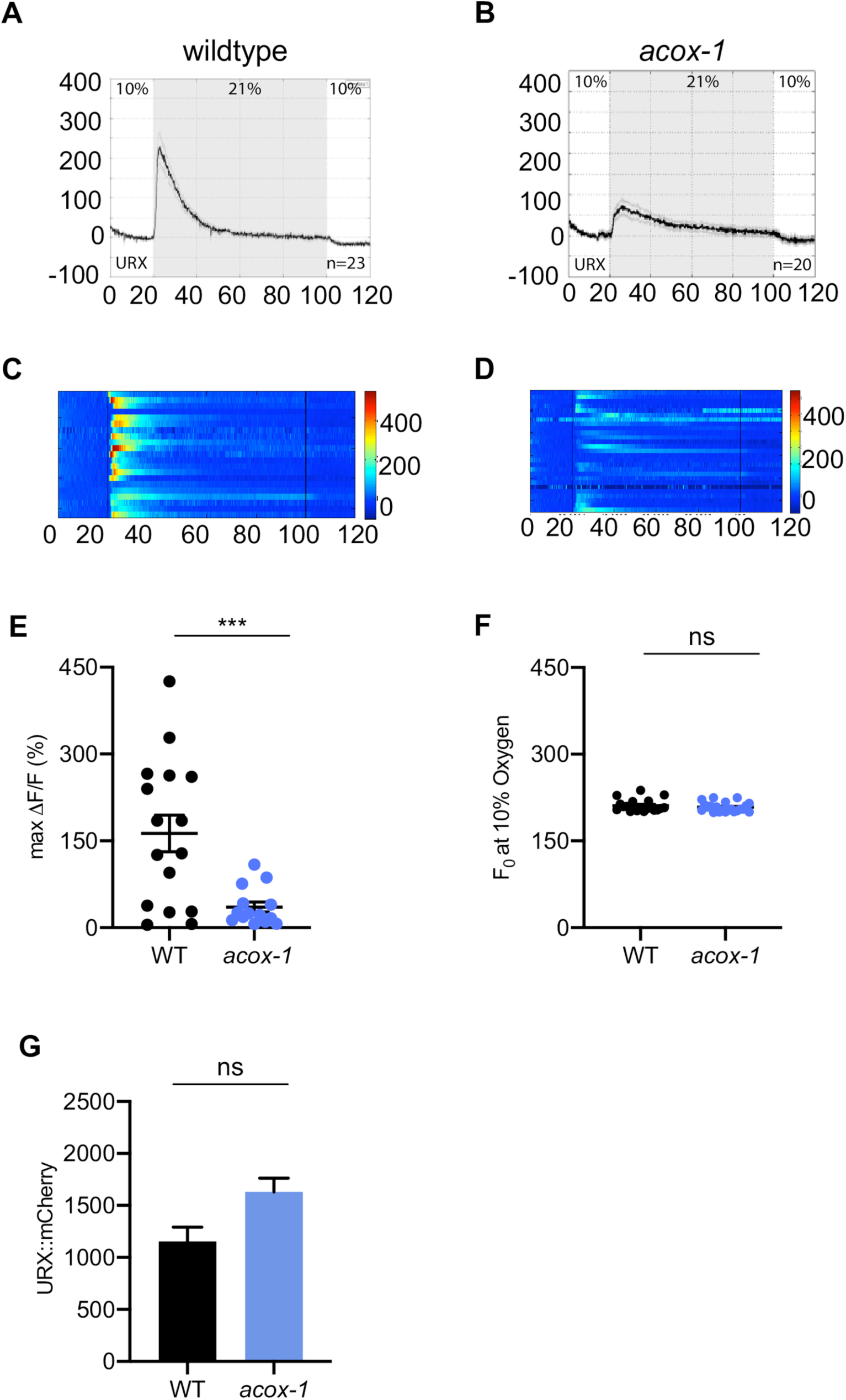
Loss of *acox-1* suppresses URX body cavity neuron activity. **(A-D)** Activity of URX neurons in each indicated genotype using Ca^2+^ imaging by GCaMP5K under the control of the URX specific *flp-8* promoter. Oxygen concentrations in the microfluidic chamber were 10% and 21% as indicated. **(A-B)** For each genotype, black traces show the average percent change of GCaMP5k fluorescence (FΔ/F_0_) and gray shading indicates SEM. The number of animals used for each condition is shown in the figure. **(C-D)** Individual URX responses are shown for each genotype; each row represents one animal. **(E)** Maximal (FΔ/F_0_) values are shown for individual animals in wildtype and *acox-1* animals. Bars indicate the average value within each genotype. ***p<0.001 by students t-test. **(F)** Individual baseline fluorescence (F_0_) values at 10% oxygen are shown for individual animals in wildtype and *acox-1* mutants. Bars indicate the median value within each genotype; n.s., not significant by students t-test **(G)** We imaged mCherry fluorescence in wildtype and *acox*-1 mutant animals expressing both GCaMP5K and mCherry under the control of the *flp-8* promoter. Images were taken in animals exposed to 10% oxygen. For each genotype, the fluorescence intensity was imaged at the same exposure, determined to be within the linear range. Fluorescence intensity was quantified and is expressed as the mean +/− SEM (n = 23). n.s., not significant by students t-test.

Our data supports a model in which loss of ACOX-1 leads to the generation of a metabolic signal derived from a fatty acyl-CoA that regulates the activity of the URX body cavity neuron in an EGL-2 dependent manner. In turn, this circuit modulates the excitability of serotonergic circuits to link internal metabolic information with behavioral responses.

## Discussion

Animals adopt distinct behavioral and physiological states in response to changes in internal metabolic status. In this study, we show that peripherally generated signals act to modulate neurally regulated processes in *C. elegans*. Specifically, we found that loss of an intestinal peroxisomal acyl-CoA oxidase leads to the production of interoceptive signals of metabolic status that modulate serotonergic regulation of feeding and egg laying. These signals are generated when acyl-CoAs that would normally be destined for peroxisomal β-oxidation via ACOX-1 are redirected into other pathways resulting in a vast change to the animal’s metabolome despite a very modest transcriptional effect. Moreover, we found that the metabolic changes in the periphery affect the activity of a specific neuron, URX, a ring neuron that is anatomically well positioned to sense peripheral signals. A change in the activity of URX dampens the effects of elevated serotonin signaling on feeding and egg laying. The precise mechanisms by which URX neurons intersect with serotonergic circuits of feeding and egg-laying remain to be determined.

Our genetic and pharmacological analyses suggest that the feeding regulatory response begins with an accumulation of a fatty acyl-CoA species. Physiological condition in which generation of acyl-CoAs exceeds their utilization predicted to occur during periods of nutrient excess, thus the feeding reducing effects of these signals are consistent with satiety-like signals. In mammals, pharmacologic or dietary manipulations that lead to elevated circulating fatty acyl-CoA levels inhibit food intake [111,112]. Fatty acyl-CoAs are short-lived species that are substrates for numerous enzymatic pathways and it is unclear in mammals whether the anorectic effect is mediated directly by specific fatty acyl-CoAs or downstream metabolic derivatives [113]. While we could not measure an obvious increase in acyl-CoA pools extracted from *acox-1* mutants, our metabolomic analyses revealed many hundreds of differentially expressed features in *acox-1* mutants, the majority of which remain structurally unidentified. The enormous metabolomic change made it unrealistic for us to pinpoint the precise metabolic species that underlies the effects of loss of *acox-1* on serotonergic signaling. Nevertheless, our metabolomics analyses revealed several classes of compounds including a variety of phospholipid species that may underlie the noted behavioral changes. Several *N*-acylethanolamines have well-known effects on mammalian feeding behavior and mood [78,114–117]. Their identification here highlighting the deep evolutionary origins of the links between metabolic state and neural mechanism that influence behavior.

We found that body cavity neurons, most prominently URX, serve as key components of the sensory circuit linking peripheral metabolic information with feeding behavior. Body cavity neurons are known to regulate oxygen sensing and social aggregation behaviors though to our knowledge they have not been implicated in regulation of pharyngeal pumping [108,110,118,119]. Given their unique anatomical placement, these neurons have long been hypothesized to facilitate bi-directional communication between the nervous system and peripheral tissues, particularly in the context of energy homeostasis. Several lines of evidence, in addition to our findings, offer experimental credence to this hypothesis. First, these neurons are in direct synaptic contact with environment sensing neurons like ADF and release modulatory signals to influence peripheral processes such as body growth, lifespan control and lipid metabolism [102,106,120,121]. Second, these neurons have the capacity to sense internal metabolic cues and integrate this information with external cues of nutrient availability to orchestrate cohesive and context-appropriate physiological responses [102]. Lastly, our data and a recent study suggest that internal nutrient sensing pathways modulate the activity of these neurons to regulate nutrient-dependent behavioral states [122]. Skora *et al.* suggest that prolonged nutrient deprivation increases the activity of URX neurons to control starvation-induced quiescence behaviors in a mechanism that involves DAF-2/IGF signaling. This data complements our findings and suggests that states of nutrient excess and nutrient deprivation have inverse effects on URX activity.

An important future challenge is to determine how the vast metabolic changes in the periphery are sensed by the body cavity neurons including URX. In principle, it is possible that some of the peripheral metabolites leave their intestinal sites of generation to directly act on the URX neurons. Alternatively, it is possible that an endocrine response originating in the intestinal cells communicates the metabolic status of the intestine to the URX neurons. Regardless of which model may ultimately be valid, our findings point to *egl-2* as a modulator of URX activity. EGL-2 is a *C. elegans* potassium channel with close homology to human ether-a-go-go related channels [98,123]. While mammalian EAG channels have not yet been implicated in the control of nutrient-dependent behavioral plasticity, numerous potassium channels have well-documented roles in metabolic sensing and energy homeostasis. For example, hypothalamic Kir6.2 K_ATP_ channels are responsive to circulating glucose levels and regulate appetite and glucose homeostasis [124–126]. Ketogenic diets, which increase circulating ketone body levels, have been shown to suppress the excitability of GABAergic neurons and reduce seizure susceptibility in a K_ATP_ channel dependent manner [127–129]. Numerous metabolites including phospholipids, polyunsaturated fatty acids, eicosanoids, fatty acyl-CoAs and oxygen can directly bind certain potassium channels to modulate channel activity [130–132]. Alternatively, metabolic information can be coupled to potassium channels by indirect signaling events involving second messengers like Ca^2+^ and cAMP, neurotransmitters or post-transcriptional modifications [133,134].

In both mammals and *C. elegans*, elevating serotonin signaling is associated with fat loss [18,33,34,135]. In mammals, this has been primarily attributed to the anorectic effects of serotonin. By contrast, detailed analysis of serotonergic effects in *C. elegans*, revealed that distinct molecular mechanisms underlie the fat and feeding effects of serotonin and that serotonin induced fat reduction in *C. elegans* is primarily driven by upregulation of peripheral mechanisms of triglyceride lipolysis and β-oxidation [18,31,32]. Although it is general overshadowed by the feeding behavioral data, the existing data indicate that serotonin also causes an increase in metabolic rate and increase fat oxidation in mammals [30,136–138]. In this study, we found that if β-oxidation pathways cannot utilize the influx of acyl-CoAs that are generated upon mobilizing stored triglycerides, a homeostatic signal is generated to blunt serotonergic effects, including effects on feeding. The complexity of the mammalian fat and feeding pathways has made it difficult to pinpoint the precise mechanisms through which serotonin levels act as satiety signals. Our findings raise the possibility that in both worms and mammals, increasing serotonin signaling may result in peripheral metabolic changes that, in turn, feedback to the nervous system to modulate food intake.

## Materials and Methods

### Worm Strains and General Maintenance

*C. elegans* strains were cultured under standard growth conditions [139]. The Bristol N2 strain was used as wildtype in all experiments and the following mutant alleles and transgenic strains were analyzed: *acox-1(ok2257), aak-2(ok524), egl-2(rg4), egl-2(n2656), daf-22(m130)*, CX10386 *kyEx2491*[*gcy-36::pkc-1*(gf)::SL2::GFP; *ofm-1::dsRed*], SSR1070 *flp-8::mCherry; flp-8::GCaMP5k*. When generating double mutants, genotypes were confirmed by PCR or sequencing. Unless stated otherwise, animals were cultured on NGM agar plates with OP50 *E. coli* at 20°C. For all experiments, worms were plated as synchronized L1 populations after hypochlorite treatment of gravid adults.

### Plasmid construction and transgenesis

Plasmids were constructed using Gateway Cloning Technology (Life Technologies). Promoter regions were amplified from wildtype genomic DNA using Phusion DNA polymerase (New England Biolabs) and sub-cloned into the pDONR-P4-P1R Gateway entry vector by BP recombination. Unless otherwise specified, primers pairs were designed based on Promoterome recommendations. For *acox-1* tissue specific rescue lines, full *F08A8.1* genomic coding sequence was amplified from wildtype genomic DNA was subsequently sub-cloned into the P221 Gateway entry vector by BP recombination. Transgenic animals were generated by injecting purified plasmids into the gonads of wildtype or mutant animals. Transgenes and an *unc-122*::*gfp* co-injection marker were injected at a concentration of 50 ng/μl. At least two independent and stably expressing lines were maintained and analyzed.

### Serotonin, Fluoxetine, Oleic Acid and Triacsin C Treatments

Stock solutions of serotonin hydrochloride (TCI America, S0370) and fluoxetine hydrochloride (Matrix Scientific, 047891) were prepared in water. For feeding experiments, synchronized L1 animals were grown on OP50 plates containing 5mM serotonin and assayed at the day 1 adult stage. To determine egg-laying responses, animals were exposed to 0.5mg/mL fluoxetine in M9 buffer or 5mg/mL serotonin in M9 buffer for 20 minutes prior to counting released eggs. Oleic acid and Triacsin C treatments were conducted as previously described [18]. Briefly, Oleic Acid (Sigma-Aldrich, O1383) was solubilized in 45% (w/v in dH_2_O) 2-hydroxypropyl-ß-cyclodextrin (Sigma-Aldrich, H5784) to 1M, and then added to OP50 plates to a final concentration of 1µM. Triacsin C (Enzo Life Sciences, BML-EI218) was solubilized in DMSO and used at 1µM on OP50 plates.

### Pharyngeal Pumping

Contractions of the posterior pharyngeal bulb were counted during a 10 second interval as previously described [18]. For measurements on fasted then refed animals, ad-libitum fed day 1 adults were washed 3 times in S-basal buffer to remove residual E. coli and subsequently placed on sterile NGM plates. Animals were fasted on plate for the indicated time then transferred to either vehicle (OP50 *E. coli*) or treatment (5mM serotonin + OP50) containing plates for 90 minutes to assay post-fast feeding responses.

### RNAi Treatment

Overnight cultures of HT115 *E. coli* containing RNAi plasmids were induced with 6mM IPTG for 4 hours at 37°C. Cultures were concentrated 2X and added to RNAi plates containing IPTG, carbenicillin, and tetracyclin. Synchronized L1 animals were added to plates and grown for 3 days at 20°C and assayed as day 1 adults.

### Microscopy

Animals were mounted on 2% agarose pads, paralyzed with NaN_3_ and imaged using a Zeiss Axioplan 2 microscope with a 16X (0.55 NA) oil immersion objective. DIC images of eggs retained *in utero* were acquired from animals at the Day 1 adult stage.

### Acyl-CoA Quantification

We adapted an HPLC-based acyl-CoA extraction, derivatization and quantification protocol from Larson et al., 2008 [61]. Briefly, 15,000 synchronized L1s were grown in liquid S-medium culture at 20°C on a rotary shaker. Animals were grown to the L4 stage and washed 3X in S-Basal then snap frozen in liquid nitrogen and stored at −80°C until further processing. To prepare lysates, pellets were thawed on ice and resuspended in 300µL of freshly prepared extraction buffer (2mL 2-propanol, 2mL pH 7.2 50mM KH_2_PO_4_, 50µL glacial acetic acid, 80µL 50mg/ml BSA). ∼500 mg of zirconium oxide beads (NextAdvance, 0.5 mm diameter, 5.5 g/ml) were added to each sample and animals were lysed using a bead beater (5 cycles - 30s ON, 1 min OFF) at 4 °C. Lysates were separated from beads by pipet and an aliquot was preserved for protein quantification (Bio-Rad, DC Protein Assay). Lysates were then washed of lipids and pigments in 200µL petroleum ether (40-60°C) saturated 1:1 with 2 propanol:water. 5µL of saturated (NH_4_)_2_SO_4_ was added to samples before extracting acyl-CoAs with 600µL 2:1 methanol:chloroform. Samples were vortexed and incubated at room temperature for 30 minutes before centrifugation. Supernatants were transferred to glass tubes and dried at 40°C in a GeneVac (∼2hrs). Once dry, samples were reconstituted in 55µL of chloroacetaldehyde derivatizing reagent (0.15 M sodium citrate, 0.5% SDS (w/v), 0.5 M chloroacetaldehyde, pH 4) and incubated in an 80°C water bath for 20 minutes. Samples were again clarified by centrifugation, before being transferred to HPLC sample tubes. 20µL of each sample was injected into an equilibrated C18 reversed-phase HPLC column (Phenomenex Luna, 4.6 mm × 150 mm, 5 µm silica particle, 100 Å pore size). A linear gradient of 0.25% (v/v) triethylamine in water and 90% (v/v) acetonitrile in water at a 0.5mL/min flow rate was used to elute acyl-CoAs. Full elution protocol described in [61]. Derivatized acyl-CoAs were detected using a fluorimeter with flash rate 100Hz and with excitation wavelength at 230nM, emission wavelength at 420nM and slid width at 20nM. Peak areas were integrated and quantified using Agilent ChemStation software.

### RNA-Seq Sample Preparation

Total RNA was extracted from ∼10,000 synchronized L4 animals using the standard Trizol, chloroform, isopropanol protocol with on-column DNAse digest. Sample quality and quantity were assessed on the Agilent Bioanalyzer using RNA 600 Nano chips and only samples with RIN score ≥9 were used for library construction. mRNA was enriched from 1µg of total RNA using NEXTflex™ Poly(A) Beads (Bioo Scientific, NOVA-512979) and strand-specific directional libraries were generated using the NEXTflex™ Rapid Directional RNA-Seq Kit (Bioo Scientific). For each genotype, samples were prepared from 3 biological replicates and indexed with distinct barcodes. Quality and fragment size distribution of synthesized cDNA libraries were assessed on the Agilent Bioanalyzer using DNA 1000 chips. Library concentrations were quantified using the NEBNext Library Quant Kit (E7630S) and each library was normalized to 10nM in TE buffer prior to pooling. Multiplexed libraries were sequenced using 100bp pair-end reads on the Illumina HiSeq 4000 platform at the UCSF Center for Advanced Technologies.

### RNA-Seq Analysis

Transcriptomic analyses were performed on the Galaxy Platform [140]. Sequencing reads were filtered and trimmed to remove barcode sequence using fastx_trimmer (http://hannonlab.cshl.edu/fastx_toolkit/). Clipped paired-end sequences were aligned to the *C. elegans* ce11 reference genome using TopHat version 2.1.1 [141]. Read counts for each gene were quantified using *htseq-count* based on gene annotations from reference annotation WS235 [142]. Differential expression analysis was conducted using the DESeq2 package (3.8) in R(3.4.1) using size factor normalization [143]. P-values were adjusted for multiple comparisons using the Benjamini-Hochberg method and a permissive false discovery threshold of q ≤ 0.1 was applied to identify differentially expressed transcripts.

### Metabolomic sample preparation

Mixed stage worms were grown in liquid and fed OP50 on days 1, 3 and 5 during the 7-day culture period, while shaking at 22 °C and 220 rpm. The cultures were centrifuged at 4 °C, and worm pellets and supernatant were frozen separately, lyophilized, and each extracted with 35 mL of 95% ethanol at room temperature for 12 h. The extracts were dried *in vacuo*, resuspended in 200µL methanol, and analyzed by LC-HRMS. All cultures were grown in at least three biological replicates.

### Mass spectrometric analysis

LC-MS analysis was performed on a Dionex 3000 UHPLC coupled with a ThermoFisher Q Exactive high-resolution mass spectrometer. Metabolites were separated using a water–acetonitrile gradient on Agilent Zorbax Eclipse XDB-C18 column (150 mm × 2.1 mm, particle size 1.8 µm) maintained at 40 °C. Solvent A: 0.1% formic acid in water; Solvent B: 0.1% formic acid in acetonitrile. The solvent gradient started at 5% B for 5 min after injection and increased linearly to 100% B at 12.5 min, and continued at 100% B for 5 min. The gradient was rapidly brought down to 5% B (over 30s) and held for 2 min for re-equilibration.

The UHPLC-MS data were collected in the profile MS mode, based on instrument specifications. Metabolites were detected as [M-H]^−^ ions or [M+Cl]^−^ adducts in the negative ionization mode (ESI+), or as [M+H]^+^ ions or [M+Na]^+^ adducts in the positive ionization mode (ESI+), using a spray voltage of 3 kV. Compound identities were confirmed based on their high-resolution masses (accuracy < 1 ppm), MS/MS fragmentation spectra, and/or comparison with authentic standards. Data analysis was carried out using the Bioconductor package XCMS [65,144]. The matched filter algorithm in XCMS for peak picking in the profile data was used.

### GCaMP Calcium Imaging

Animals were exposed to different oxygen concentration (10% or 21%) using a microfluidic chamber constructed with the oxygen-permeable poly(dimethylsiloxane) (PDMS) as described [105]. A Valvebank II (AutoMate Scientific, Inc.) was used to control input from two pressurized pre-mixtures of oxygen and nitrogen containing either 10% oxygen or 21% oxygen (Praxair, Inc.). The gas flow rate was measured by a VWRTM traceable pressure meter and set to 0.26 psi. At the time of experiment, an individual day 1 adult animal was picked without using any food and consecutively transferred to two unseeded plates immediately before imaging. The transferred animal was immobilized in S basal containing 5 mM levamisole and transported into the microfluidic chamber via Tygon tubing (Norton). To avoid drying, the animal was constantly submerged in S-Basal buffer while inside the chamber and GCaMP5K fluorescence was visualized at 40x magnification using a spinning disk confocal microscope (Olympus) with MetaMorphTM software (version 6.3r7, Molecular Devices). As described earlier, worms were pre-exposed to 10% oxygen for 5 min in the microfluidic chamber [105]. GCaMP5K fluorescence was recorded by stream acquisition for 2 min at a rate of 8.34 frames/second, with an exposure time of 20ms using a 12-bit Hamamatsu ORCA-ER digital camera. Each animal was recorded once, and GCaMP5K-expressing neurons were marked by a region of interest (ROI). The change in fluorescent intensity as per neuronal excitation and position of the ROI was tracked using the “Track Objects” function in MetaMorphTM. To subtract background from the total integrated fluorescence intensity of the ROI, an adjacent ROI was selected in the same image. MATLAB (MathWorks,Inc.) was used to analyze the data. Fluorescence intensity is presented as the percent change in fluorescence relative to the baseline (ΔF/F0). F0 was measured in worms exposed to 10% oxygen during the first 9-13 seconds for each recording and calculated as an average over that period. The number of animals used for each condition is denoted in the figures.

### Statistics

Graphpad Prism 8.0 software package was used to calculate all p-values unless otherwise specified. Where only 2 conditions were compared, a two-tailed students t-test was performed. ANOVAs with appropriate post-test corrections were used when comparing multiple conditions.

## Acknowledgements

We thank the *Caenorhabditis* Genetics Center (CGC), which is funded by the NIH Office of Research Infrastructure Programs (P40 OD010440), for providing some of the strains used in this study. We also thank the De Bono lab for sharing the *gcy-32*::*egl-2*(gf) construct and the Bargmann Lab for the generous gift of the CX10386 strain. Lastly, we thank George Lemieux for helpful advice and discussion and members of the Ashrafi lab for comments on the manuscript.

---

**Figure S1, related to Figure 1.**
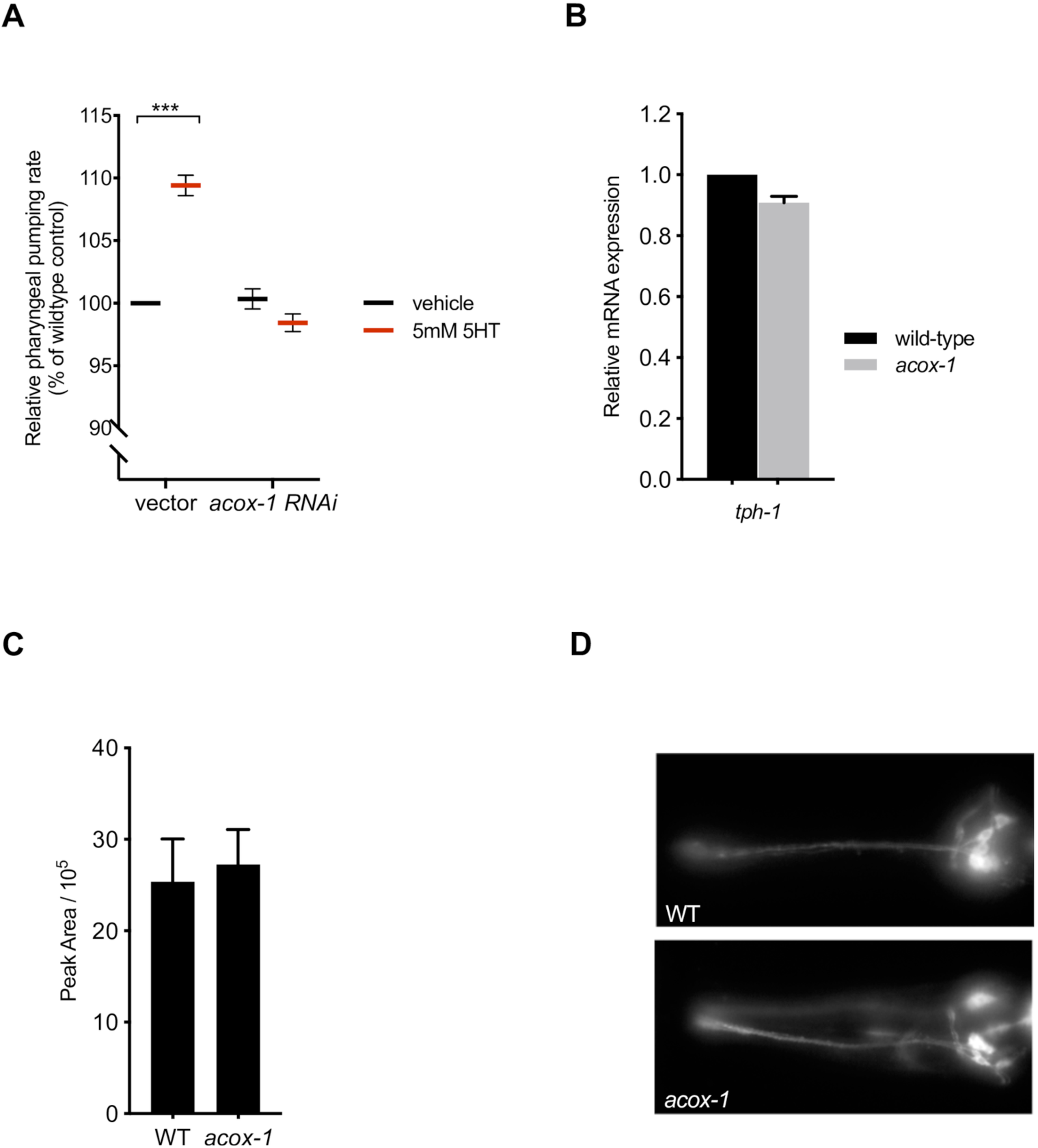
**(A)** Knocking down *acox-1(F08A8.1)* by RNAi suppresses the feeding elevating effects of exogenous serotonin (5mM 5HT). Animals were treated with RNAi from L1 stage and feeding was assayed at day 1 adult stage. Feeding data is expressed as a percentage of vehicle treated wildtype animals. Error bars indicate +/− SEM from mean, n = 20 animals per strain. *** p < 0.001 ANOVA (Tukey) **(B)** Loss of *acox-1* does not influence the transcriptional expression of tryptophan hydroxylase (*tph*-1) as measured by qPCR. Error bars indicate +/− SEM from mean n = 3 independent assays **(C)** Relative abundance of 5-HT in wildtype and *acox-1* mutants, as determined by LC-HRMS. Error bars indicate +/− SEM from mean, n = 4 independent experiments. (**D**) Loss of *acox-1* does not grossly alter amphid neuron morphology. DiI staining of amphid chemosensory neurons in wildtype and *acox-1* mutants. Images acquired at day 1 adult stage.

**Figure S2, related to Figure 3.**
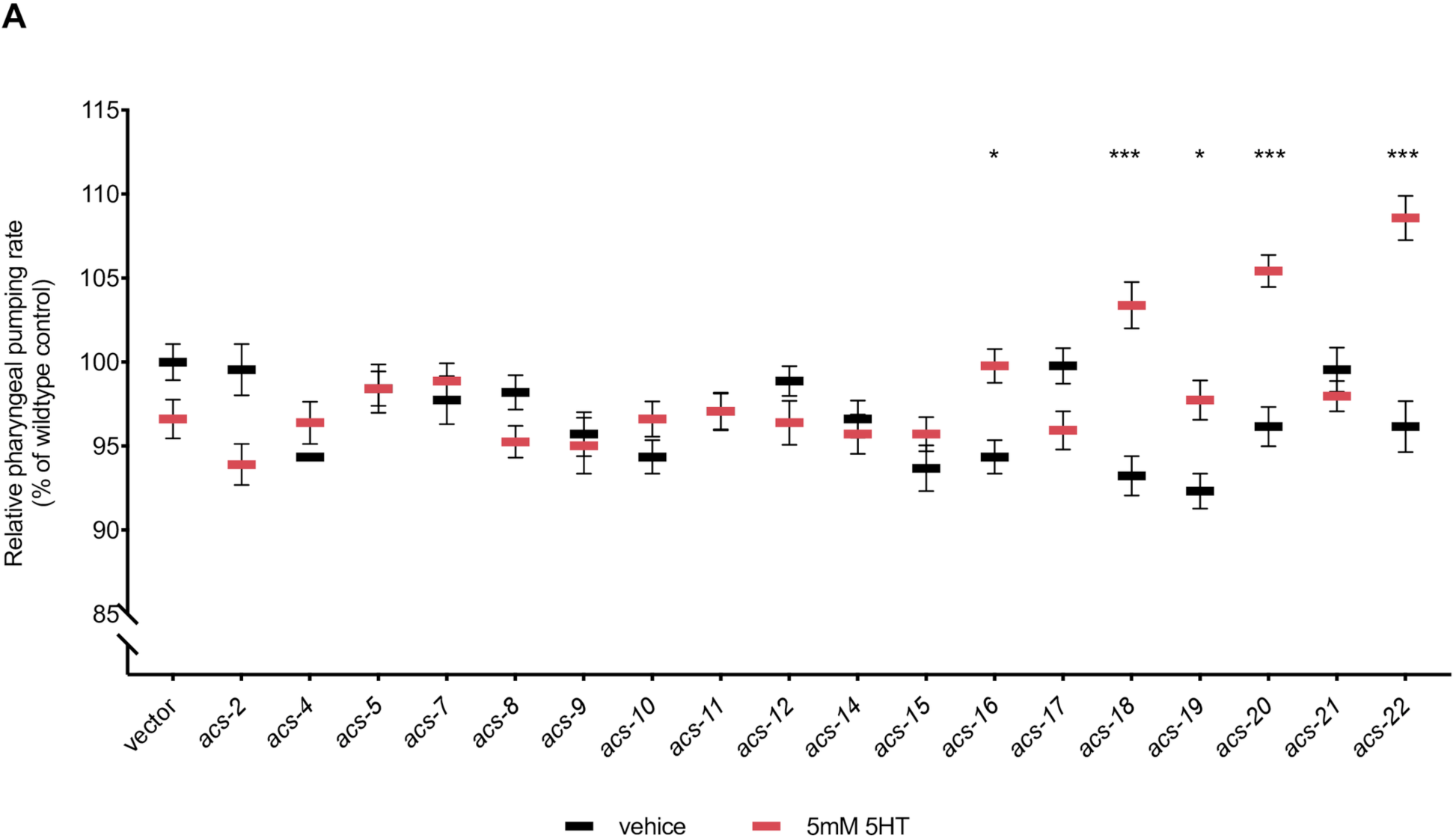
**(A)** RNAi-mediated inactivation of distinct acyl-CoA synthases suppress feeding defects in *acox-1(ok2257)* animals. Animals were treated with respective RNAi clones from L1 stage and feeding was assayed at day 1 adult stage. Feeding data is expressed as a percentage of vehicle treated wildtype animals. Error bars indicate +/− SEM from mean, n = 15 animals per strain. *<0.05, *** p < 0.001 two way ANOVA (Tukey).

**Figure S3, related to Figure 4.**
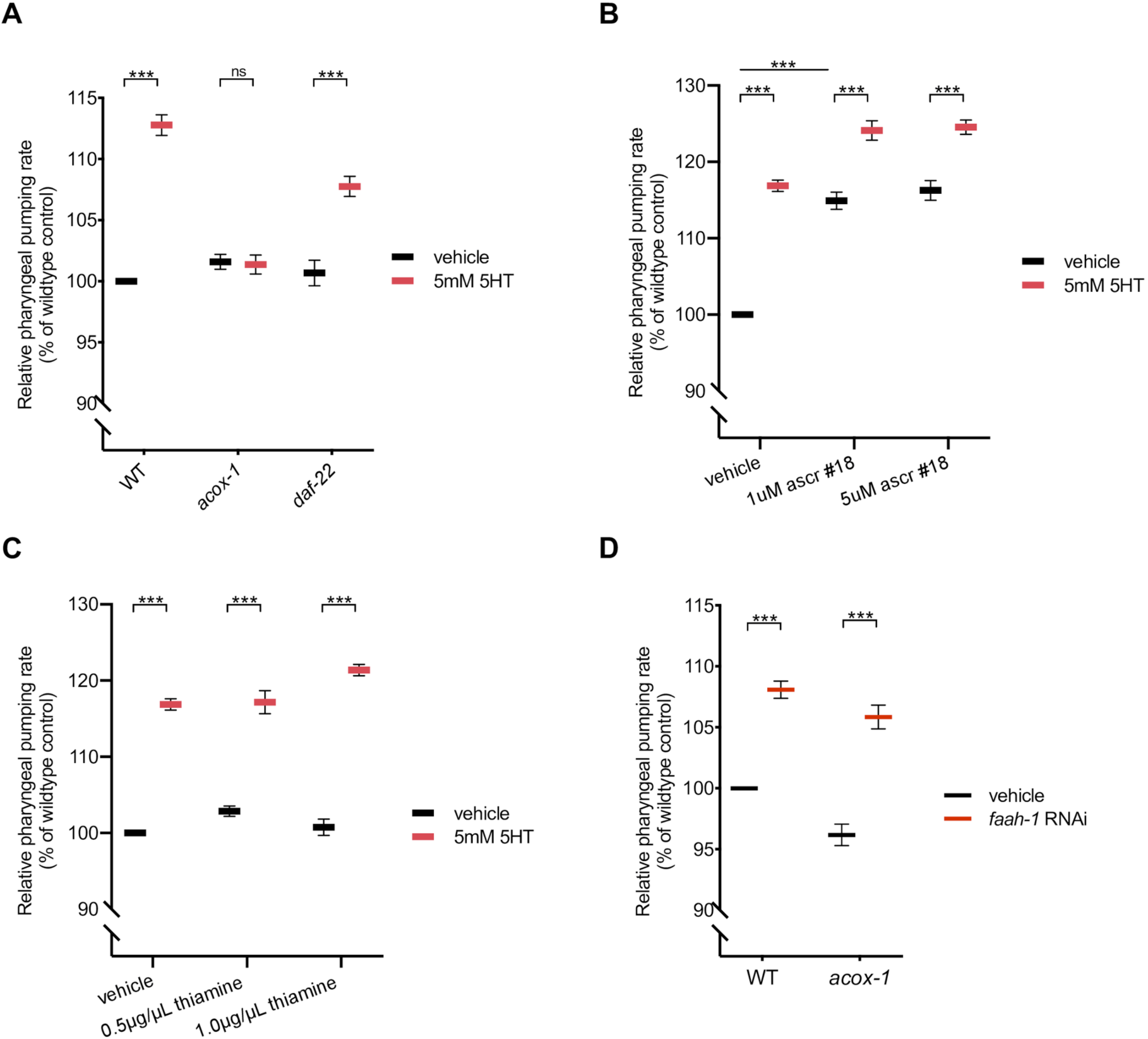
**(A)** *daf-22(ok693)* animals are still responsive to the feeding elevating effects of 5mM serotonin. n=10 animals per condition **(B)** Feeding responses of wildtype animals to ascaroside #18 (ascr #18). Animals were exposed to 1µM and 5µM ascr#18 from L1 stage and pharyngeal pumping rates were determined at day 1 adult stage. n = 15 animals per condition **(C)** Feeding responses of wildtype animals to thiamine. Animals were exposed to 0.5 µg/µL thiamine and 1.0 µg/µL thiamine from L1 stage and pharyngeal pumping rates were determined at day 1 adult stage. n = 15 animals per condition **(D)** RNAi-mediated inactivation of fatty acid amide hydrolase (*faah-1*) elevates feeding responses of wildtype and *acox-1* mutants. Animals were grown on *faah-1* RNAi from L1 stage and feeding was assayed at day 1 adult stage. n = 10 animals per condition. All feeding data is expressed as a percentage of vehicle or vector treated wildtype animals. Error bars indicate +/− SEM from mean, *** p < 0.001 two way ANOVA (Tukey).

**Figure S4, related to Figure 5.**
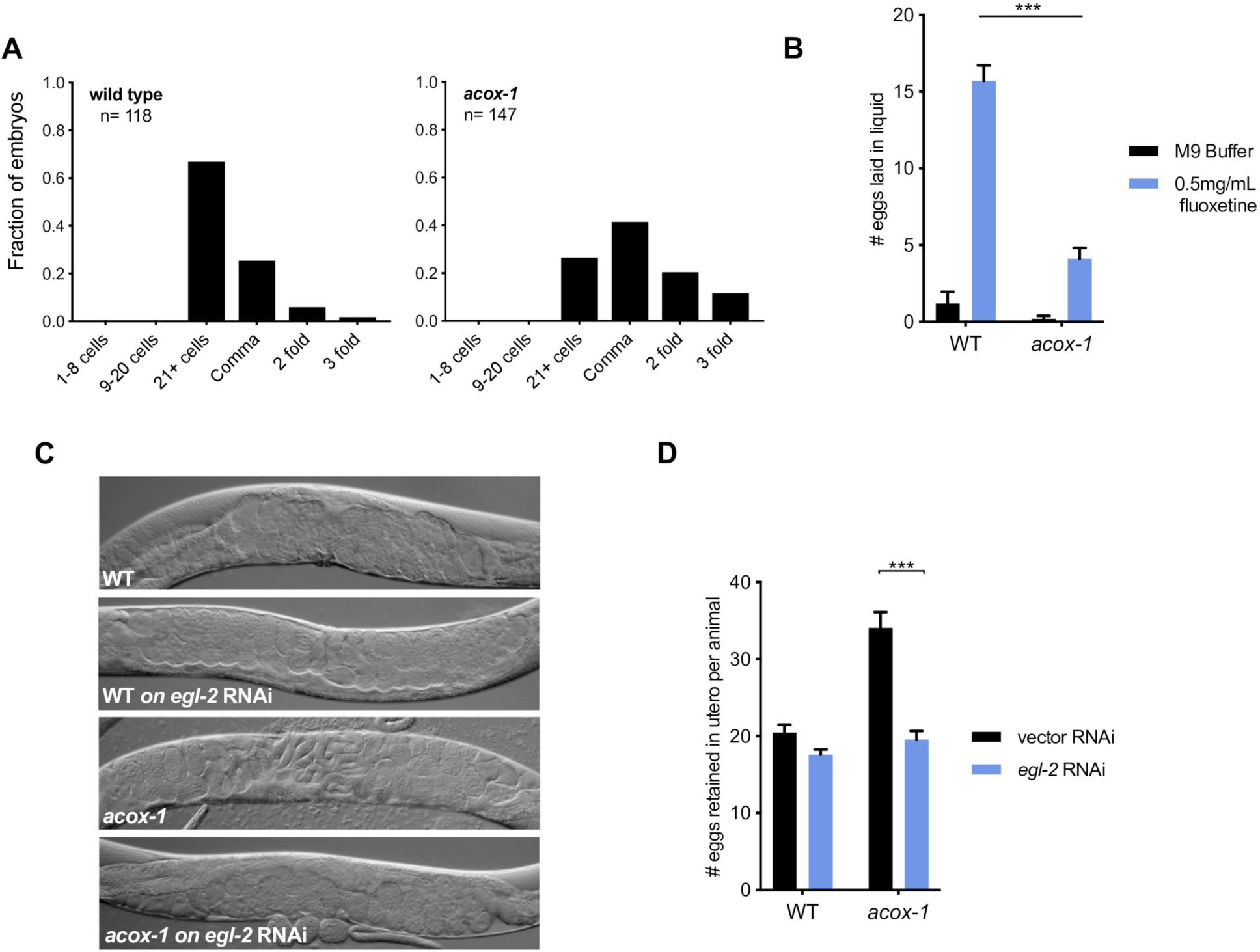
**(A)** *acox-1(ok2257)* mutants lay eggs at a later developmental stage that wildtype animals, suggesting that *in utero* retention time is increased. Histograms indicate the distribution of embryos at each developmental stage. **(B)** *acox-1* mutants are less responsive to the egg-laying inducing effects of serotonin. Egg-laying response of wildtype and *acox-1* mutants in control buffer (M9) or 0.5mg/mL fluoxetine. Data represents the number of eggs released per animal after a 20-minute exposure to vehicle or drug. Error bars represent +/− SEM from mean. n = 15 animals per condition, *** p < 0.001 ANOVA (Sidak) **(C-D)** Inactivation of *egl-2* via RNAi rescues *acox-1* egg-laying defects. Representative DIC images of day 1 adults of each genotype **(C)** and quantification **(D)** of eggs retained *in utero*. Error bars indicate +/− SEM from mean, n = 15 animals per genotype. *** p < 0.001 unpaired student’s t-test.

**Figure S5, related to Figure 6.**
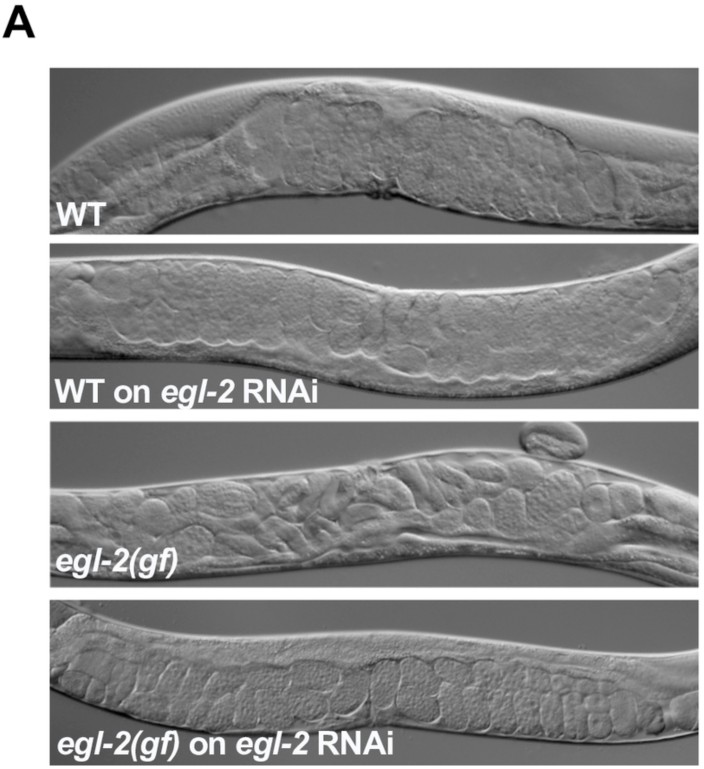
**(A)** RNAi-mediated knockdown of *egl-2* rescues egg-laying defects associated with aberrant channel activity *egl-2(n698)* mutants. Representative DIC images acquired from day 1 adults.

